# A morpho-transcriptomic map of brassinosteroid action in the Arabidopsis root

**DOI:** 10.1101/2021.03.30.437656

**Authors:** Moritz Graeff, Surbhi Rana, Jos R. Wendrich, Julien Dorier, Thomas Eekhout, Ana Cecilia Aliaga Fandino, Nicolas Guex, George W. Bassel, Bert De Rybel, Christian S. Hardtke

**Affiliations:** Department of Plant Molecular Biology, University of Lausanne, Biophore Building, 1015 Lausanne, Switzerland; Ghent University, Department of Plant Biotechnology and Bioinformatics, Technologiepark 71, 9000 Ghent, Belgium; VIB Center for Plant Systems Biology, Technologiepark 71, 9000 Ghent, Belgium; Bioinformatics Competence Center, University of Lausanne, Genopode Building, 1015 Lausanne, Switzerland; School of Life Sciences, The University of Warwick, Coventry, CV4 7AL, UK

**Keywords:** Arabidopsis, root, meristem, brassinosteroid, single-cell mRNA sequencing, segmentation, morphology, BRI1, digital single-cell analysis

## Abstract

The effects of brassinosteroid signaling on shoot and root development have been characterized in great detail but did not identify a simple consistent positive or negative impact on a basic cellular parameter that would comprehensively explain the phenotype of brassinosteroid-related mutants. Here we combined digital 3D single-cell shape analysis and single-cell mRNA sequencing to characterize root meristems and mature root segments of brassinosteroid-blind mutants and wildtype. These data demonstrate that brassinosteroid signaling neither affects cell volume nor cell proliferation capacity. Instead, brassinosteroid signaling is essential for the precise orientation of cell division planes and the extent and timing of anisotropic cell expansion. Moreover, we found that the cell-aligning effects of brassinosteroid signaling can propagate to normalize the anatomy of both adjacent and distant brassinosteroid-blind cells through non-cell-autonomous functions, which are sufficient to restore overall root growth vigor. Finally, single-cell transcriptome data discern directly brassinosteroid-responsive genes from genes that can react to non-cell-autonomous brassinosteroid-dependent signals and highlight arabinogalactans as sentinels of brassinosteroid-dependent anisotropic cell expansion.

## INTRODUCTION

Steroids are found throughout the tree of life and have acquired hormone function in animals as well as in plants (Ferreira-Guerra et al., 2020; Tarkowska, 2019). The brassinosteroids comprise a few bioactive molecules, among which brassinolide is typically the natural, endogenous ligand in dicotyledons (Nomura et al., 2005; Roh et al., 2020). In the model plant *Arabidopsis thaliana* (Arabidopsis), brassinolide is sensed by the extra-cellular domains of the receptor kinase BRASSINOSTEROID-INSENSITIVE 1 (BRI1) and its close homologs, BRI1-LIKE 1 (BRL1) and BRL3 (Cano-Delgado et al., 2004; Li and Chory, 1997). Their interaction with brassinolide triggers a phospho-transfer cascade, which eventually increases the nuclear abundance of downstream transcriptional effectors and thereby modulates the expression of their target genes (Kim and Russinova, 2020; Tang et al., 2016; Wang et al., 2002; Yin et al., 2002). Among them, brassinosteroid biosynthesis pathway genes are prominent, which establishes a negative feedback that maintains brassinosteroid signaling homeostasis (Wang et al., 2002). Loss-of-function mutations in genes encoding brassinosteroid receptors, biosynthetic enzymes or downstream effectors lead to severe growth retardation, which can vary in its extent depending on allele strength and redundancies (Cano-Delgado et al., 2004; Chen et al., 2019; Choe et al., 1998; Li and Chory, 1997; Szekeres et al., 1996). The dwarf phenotype of brassinosteroid-related mutants has been interpreted from various angles (Oh et al., 2020). Although overall reduced cell size appears to be a consistent phenotype, effects of brassinosteroids on cell proliferation might be context-dependent. For example, whereas a brassinosteroid signaling gain-of-function slightly promotes cell proliferation in Arabidopsis leaves (Gonzalez et al., 2010; Vanhaeren et al., 2014), it represses cell proliferation in rice leaf lamina (Sun et al., 2015). Yet detailed analysis of a biosynthesis mutant suggests that brassinosteroids promote the exit from cell division, and thereby also the transition to cell expansion and differentiation (Zhiponova et al., 2013). Moreover, it has been proposed that brassinosteroid signaling is required for the differentiation of vascular tissues (Cano-Delgado et al., 2004; Holzwart et al., 2018; Kang et al., 2017; Tamaki et al., 2020; Yamamoto et al., 2001).

Leaves are initiated from founder cells at the flanks of shoot apical meristems. The size of the founder cell pool largely determines final leaf size, which is remarkably well buffered against fluctuation in cell proliferation or expansion by compensatory mechanisms (Hisanaga et al., 2015; Vercruysse et al., 2020). By contrast, root growth is essentially continuous and driven by an apical stem cell niche (SCN) in the root meristem tip, which is maintained by the so-called quiescent center (QC). Proximal to the SCN, daughter cells undergo repeated meristematic as well as a few highly controlled formative divisions before they eventually expand and differentiate (Figure S1A). These divisions thus give rise to precisely aligned cell files that differentiate into the individual root tissues in a stereotypic pattern of overall radial symmetry, and bilateral symmetry in the vascular cylinder (Figure S1B). Finally, the timing of differentiation is not uniform across cell files and tissues. Thus, although the overall shape and organization of the root meristem is more recognizable and stereotypic than in the shoot meristem, cell proliferation, elongation and differentiation are by comparison more intimately intertwined.

The role of brassinosteroid signaling in root development appears to be complex (Singh and Savaldi-Goldstein, 2015). Opposing, action-site-dependent effects on cellular differentiation have been reported (Vragovic et al., 2015), and with regard to cell proliferation, interpretations of phenotypes vary. For example, root meristem size in *bri1* mutants could be reduced as indicated by the number of meristematic cells in cortex or epidermal cell files (Gonzalez-Garcia et al., 2011; Hacham et al., 2011), yet such reduction was not observed in a brassinosteroid biosynthesis mutant (Chaiwanon and Wang, 2015). Moreover, brassinosteroid signaling appears to restrict formative divisions (Kang et al., 2017). Cell proliferation defects together with generally decreased cell elongation could explain the strongly reduced root growth vigor in brassinosteroid mutants, and it has been proposed that brassinosteroid signaling promotes cell cycle progression (Gonzalez-Garcia et al., 2011; Hacham et al., 2011) and sets the absolute cell size in a “sizer” mechanism of cell elongation (Pavelescu et al., 2018). Indeed, the maximum elongation rate in *bri1* mutants seems to be reduced (Hacham et al., 2011) whereas the average time cells spend elongating remains similar to wildtype (Cole et al., 2014). Moreover, brassinosteroid signaling can at least in part act non-cell-autonomously, as initially proposed from the observation that *BRI1* expression restricted to the epidermis could largely rescue the *bri1* shoot growth defect (Savaldi-Goldstein et al., 2007). Restricted epidermal *BRI1* also appears to promote both cell proliferation and elongation in roots (Hacham et al., 2011), and this effect was exaggerated in *bri1 brl1 brl3* triple receptor null mutant background (*“bri^TRIPLE^”* mutants) (Vragovic et al., 2015). The observation that overall root growth is only partially rescued in such lines (Kang et al., 2017) might reflect a trade-off in the balance between cell proliferation and cell growth, which generally determines overall root meristem size and growth vigor (Svolacchia et al., 2020). However, restoration of brassinosteroid signaling exclusively in the two developing protophloem sieve element cell files of *bri^TRIPLE^* mutant root meristems, the first proximal tissue to differentiate (Furuta et al., 2014; Rodriguez-Villalon et al., 2014), can rescue root growth vigor to nearly wild type levels (Kang et al., 2017), likely via non-canonical signaling outputs (Graeff et al., 2020). Thus, brassinosteroid can act non-cell-autonomously in various ways, including through a phloem-derived, non-cell-autonomous organizing signal.

In this study, we set out to map the effects of brassinosteroid signaling in the root by the characterization of *bri^TRIPLE^* mutants at single-cell resolution. Our data suggest that brassinosteroid signaling neither affects cell proliferation nor cellular growth capacity *per se*, but rather enforces accurate cell division plane orientation and cellular anisotropy (i.e. an increased ratio between longitudinal and radial cell expansion).

## RESULTS

Because brassinosteroid biosynthesis is a branched pathway (Ohnishi et al., 2006), and because brassinosteroid signaling is subject to negative feedbacks (Ruan et al., 2018; Wang et al., 2002) and can additionally act through non-canonical outputs (Caesar et al., 2011; Cho et al., 2014; Holzwart et al., 2018; Kondo et al., 2014; Tamaki et al., 2020; Wolf et al., 2014), the brassinosteroid receptor triple null mutant represents the consequences of absent brassinosteroid action most stringently. We therefore concentrated our efforts on the *bri^TRIPLE^* mutant and contrasted it against wildtype and *bri^TRIPLE^* mutants in which brassinosteroid signaling was exclusive restored in the developing protophloem sieve elements (“*bri^T-RESCUE^”*) (Graeff et al., 2020; Kang et al., 2017).

### Brassinosteroid signaling promotes anisotropy of mature root cells

Typically, effects of brassinosteroid mutants on cell size have been described in terms of cell elongation in the root meristem, with a focus on cortex cells because of their considerable size and uniformity as compared to other root tissues (Chaiwanon and Wang, 2015; Hacham et al., 2011; Kang et al., 2017; Pavelescu et al., 2018). Here we sought to determine the impact of brassinosteroid signaling on root cell growth comprehensively, in a first instance by measuring cellular parameters of mature root cells across all tissues. To preserve root anatomy, we prepared roots according to the ClearSee protocol (Kurihara et al., 2015). They were then mounted on microscopy slides in media supplemented with 0.3% agar and with distancers to prevent their physical deformation during imaging by confocal microscopy. High resolution 3D stacks were acquired of the mature parts of ten to eleven roots per genotype (Figure 1A, Movies S1-S3). Each image stack was then processed with the *PlantSeg* segmentation software (Wolny et al., 2020) to identify and label individual cells, followed by analysis in *MorphoGraphX* (Barbier de Reuille et al., 2015) to extract cellular parameters. For subsequent single-cell 3D geometric analyses, *3DCellAtlas* (Montenegro-Johnson et al., 2015) was used, which both identified cell types and measured cellular parameters in defined cellular positions along the length of the root axis. With this approach, cells could be grouped into six categories that reflected distinct cell types, with the exception of the vascular tissues that were lumped together in one category. The quantitative analysis of hundreds of cells showed that they were overall significantly shorter in *bri^TRIPLE^* than in wildtype or in *bri^T-RESCUE^* in all tissues (Figure 1B and E). At the same time, they were also significantly wider with respect to the radial or circumferential dimensions (Figure 1C and E), which likely contributed to the overall increased thickness of *bri^TRIPLE^* roots (Figure 1A). However, with very few, borderline significant exceptions, no significant differences were observed for cell volume between the three genotypes (Figure 1D and E). In summary, our high throughput data thus indicate that loss of brassinosteroid signaling results in reduced cellular anisotropy, but does not impact cell volume.

**Figure 1.**
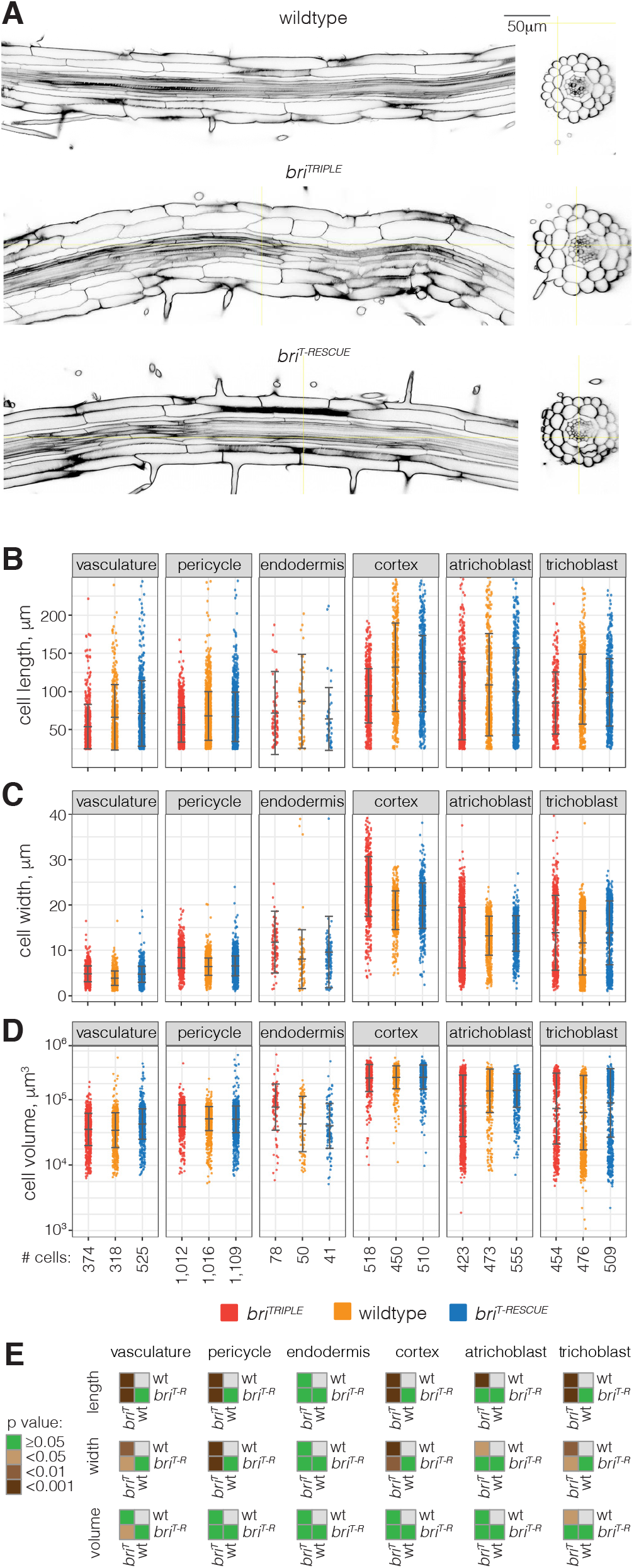
Comparison of mature wildtype, *bri^TRIPLE^* and *bri^T-RESCUE^* roots tissues. (A) Transversal and horizontal (optical) sections of representative 8-day-old primary roots, ~1 cm above the root meristem, illustrating reduced anisotropy of mature cells in *bri^TRIPLE^* roots. (B-D) Comparative quantitative analyses for different cell morphology parameters obtained by processing of 3D stacks with the *PlantSeg-MorphoGraphX-3DCellAtlas* pipeline, ten roots per genotype. Overall cell length (B) is reduced in the *bri^TRIPLE^* roots, whereas cell width (C) is increased. Average cell volume (D) is similar between the three genotypes, indicating that it is not cell expansion but rather cellular anisotropy that is reduced in *bri^TRIPLE^* mutants. Whiskers indicate mean and standard deviation. (E) Statical comparison between cell types of different genotypes (ANOVA, averages per root). Note that the lignin- and suberin-rich secondary walls of mature endodermal cells caused uneven calcofluor cell wall staining, which oversaturated some images and interfered with cell boundary prediction by the segmentation software. Obviously fused endodermis cells were discarded from the analysis. The attributes of the remaining endodermis cells might sti ll be thwarted by these difficulties, which could explain why differences in (A) and (B) were not statistically significant although the measurements display the same tendencies as in the other tissues.

### Brassinosteroid signaling does not affect cell division capacity of the root meristem

The fact that the reduction in *bri^TRIPLE^* root growth vigor cannot be explained by reduced cell elongation alone has been established previously, and additive effects from reduced cell proliferation as well as delays due to supernumerary formative divisions were deemed responsible for the extent of the macroscopic root phenotype (Gonzalez-Garcia et al., 2011; Hacham et al., 2011; Kang et al., 2017). Indeed, the estimated number of cells in 7-day-old *bri^TRIPLE^* roots appeared to be reduced (Figure S2A-B). We therefore extended our 3D cellular morphometric analysis to the root meristems (Figure 2A). Again, we obtained high resolution 3D stacks (Movies S4-S6) and processed them through the *PlantSeg-MorphoGraphX-3DCellAtlas* pipeline, with automated annotation of cell identities that were close to the SCN to assign tissue identity to individual cell files. As we were interested in the question how the growth vigor differences between the investigated genotypes arise, we concentrated our analyses on the proximal, permanent root tissues and disregarded both the columella and lateral root cap cells, which eventually undergo cell death and are shed (Fendrych et al., 2014; Xuan et al., 2016). The analysis indicated that cellular anisotropy is already reduced in meristematic *bri^TRIPLE^* cells, because close to the QC they were in general already shorter (Figure 2B) but also wider than in wildtype (Figure 2C). Moreover, *bri^TRIPLE^* cells again did not appear to be consistently smaller in terms of volume (Figure 2D). Finally, we sought to determine the number of actively dividing cells per root meristem, which can be visualized by antibody immuno-staining of cell plates by the cytokinesis-specific syntaxin KNOLLE (KN) (Lauber et al., 1997). Based on these immunostainings (Figure S2C-E), the number of actively dividing cells was indeed not significantly different between *bri^TRIPLE^* and wildtype (Figure S2F). Moreover, cell plates were in all cases (n=21-25 meristems per genotype) observed within ~200 microns distance from the QC, suggesting that the spatial window for cell divisions is neither substantially shortened nor prolonged in *bri^TRIPLE^* meristems as compared to wildtype. In summary, these analyses indicate that cells in *bri^TRIPLE^* meristems do not display a generic reduction in their proliferation capacity.

**Figure 2.**
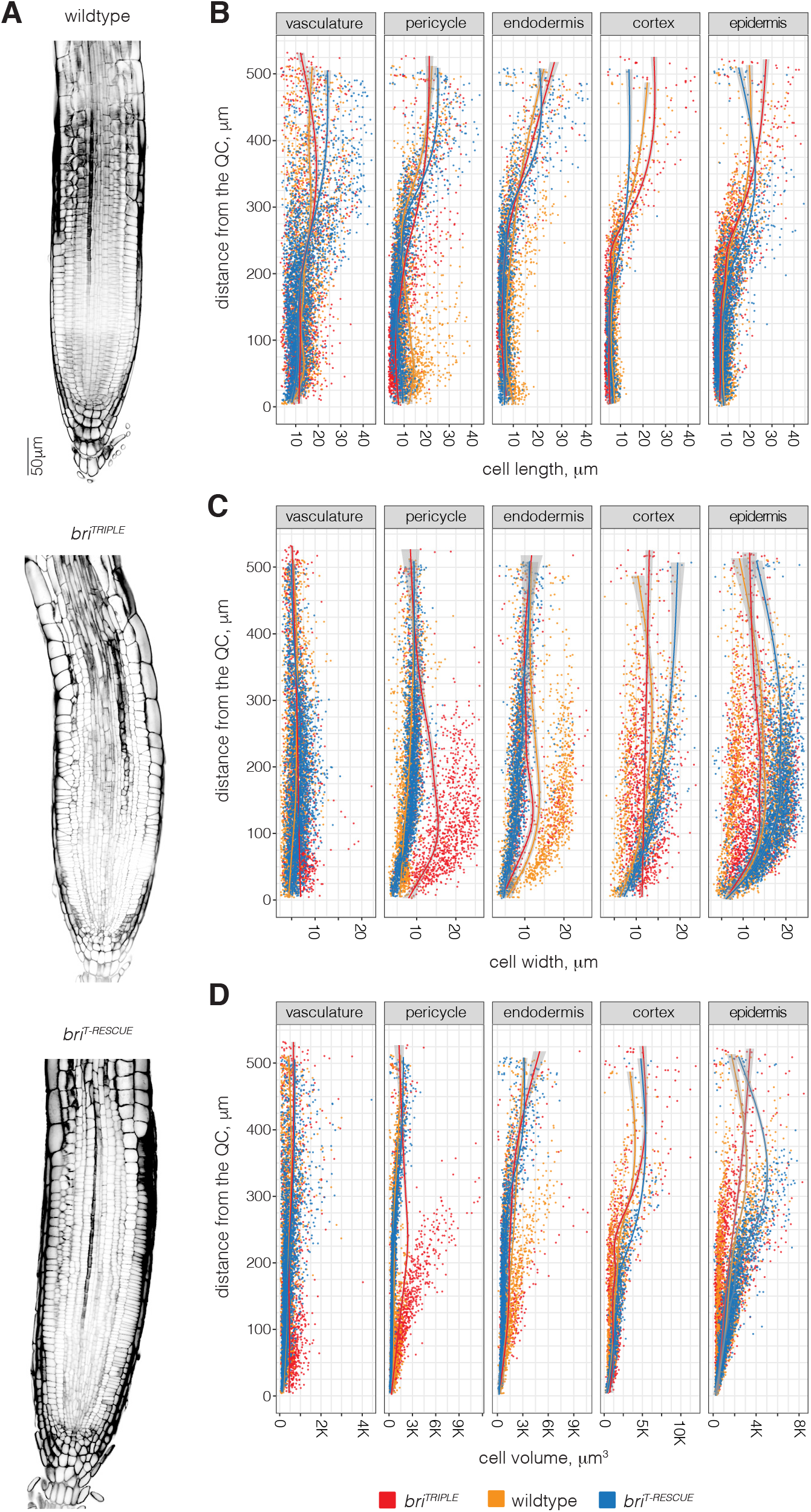
Comparison of wildtype, *bri^TRIPLE^* and *bri^T-RESCUE^* root meristems. (A) Transversal sections of representative 8-day-old primary root meristems of the three genotypes. Note the comparatively disorganized morphology of *bri^TRIPLE^* meristems. (B-D) Scatterplots of cell length (B), width (C) and volume (D) for the root meristem cell types analyzed through the *PlantSeg-MorphoGraphX-3DCellAtlas* pipeline. Models were fitted to the measurements for each genotype using a general additive model with a shrinking cubic regression spline interpolation, grey areas indicate the standard error. Variation within roots of the same genotype is generally high, making comparisons of the cellular properties difficult.

### Brassinosteroid signaling guides the transition to cell differentiation

Our initial cellular morphometric analyses produced statistically robust outputs per root for the mature parts (Figure 1B-E). However, for the root meristems, we sometimes observed substantial variation across and within genotypes between individual samples (Figure S3A-C). Such effects were compounded with increasing distance from the QC, where morphology became more variable because root meristems are dynamic, growing structures (Taylor et al., 2021; Thompson and Holbrook, 2004). Their developmental plasticity was for instance evident in the fluctuation of growth orientation (Thompson and Holbrook, 2004), which led to both within-root and between-root variation in cellular parameters of the same tissue. Moreover, because of frequently disorganized cell files in *bri^TRIPLE^* meristems (Figure 2A), positional attributes were often distorted and thus difficult to compare between roots. Therefore, we devised a method to permit more fine-grained, standardized comparisons across several roots and genotypes. In brief, we first manually marked the QC, the central xylem pole and the orientation of the xylem axis in each segmentation to define its biological main axes. These 3D meshes were then realigned by straightening of the main axis, by restoring the overall radial symmetry and by aligning the main axis with the z-axis, and the xylem axis with the x-axis (Figure 3A-C). The y-axis, perpendicular to the x-axis, consequently passed approximately through the two phloem poles. Importantly, the original cellular parameters obtained from *MorphoGraphX* were not altered by these realignments. Altogether, these transformations gave rise to idealized representations of the samples (Figure 3D-F), which could be directly compared due to their straight, cylindrical layout around a central z-axis, their alignment with respect to the bilateral symmetry of the stele, and the reference point set to the center of the QC.

**Figure 3.**
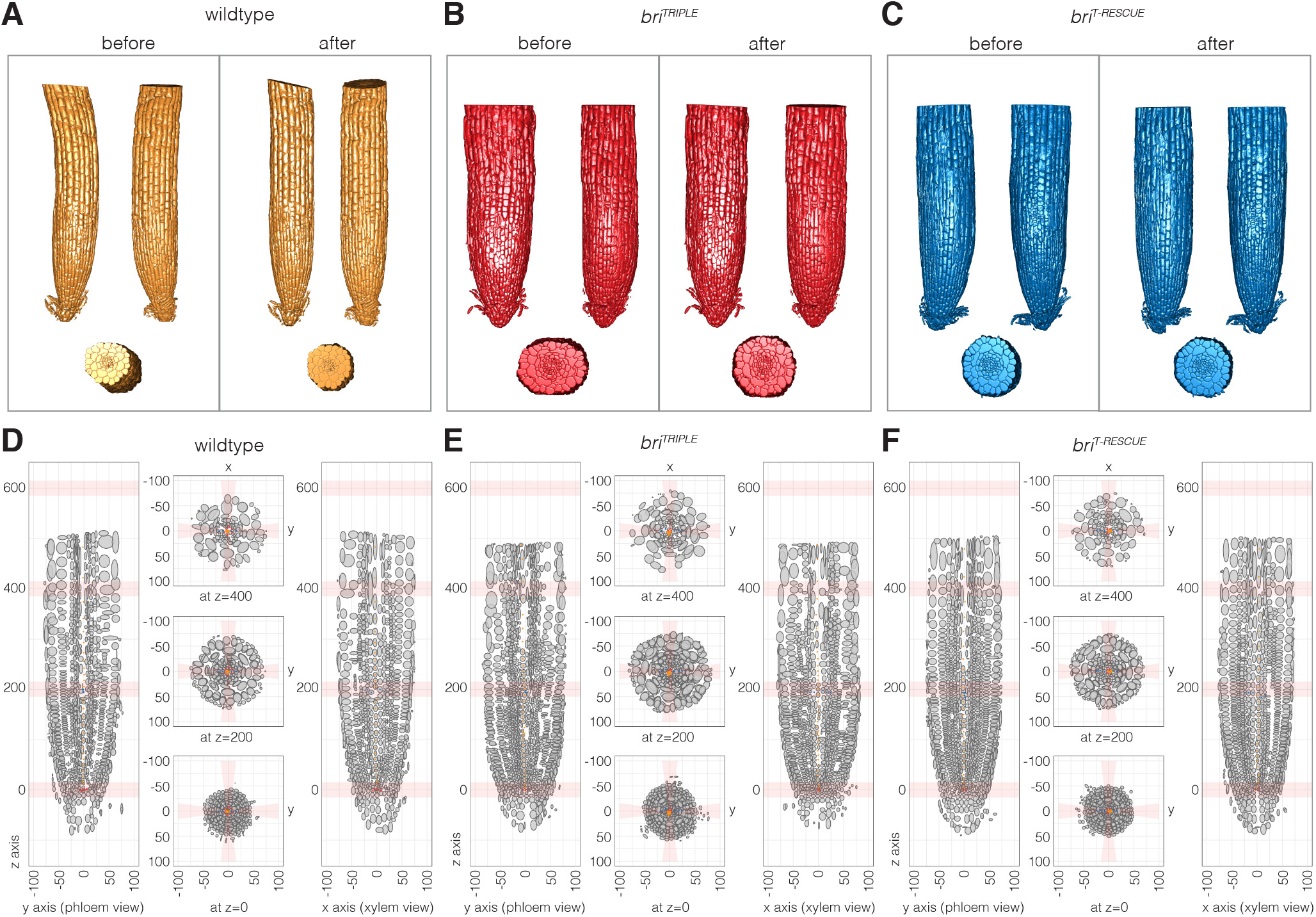
Standardization of root meristem segmentations for quantitative comparisons. (A-C) Illustration of the straightening of root meristem 3D-meshes for examples of the different genotypes, before and after realignment. (D-F) Transversal and horizontal virtual sections of simplified root meristem models obtained by re-orientation (straightening) of 3D-meshes along the defined central and radial axes. Cells are shown as ellipsoids with semi-axis proportional to the cell length in the corresponding direction. Cells of the central xylem pole are marked by orange dots, of the QC by red dots, and of the xylem axis by blue dots. For each root, the central xylem pole was re-aligned with the vertical z-axis, the xylem axis with the radial y-axis, and the phloem poles with the radial x-axis. The original cellular parameters were maintained as cells were re-oriented along the respective axis.

Our method allowed us to acquire comparable per-position parameters, and a pool of eleven to twelve roots per genotype were used for comparison of the genotypes. Unbiased quantifications were performed by radial and longitudinal sliding window analyses using concentric cylindrical shells (centered on the z-axis) of 10 micron thickness in the radial direction and 50 micron height. Analyses in the radial direction were performed by varying the radius of the shell while keeping its z-position fixed (Figures S4 and S5). These data gave a general impression of cellular features along the x- and y-axes and across tissues. They corroborated that meristematic *bri^TRIPLE^* cells were already shorter in the longitudinal dimension and wider in both the radial and circumferential dimensions close to the SCN (Figure S4A-C), whereas cell volume was comparable to wildtype (Figure S5A). For analyses in the longitudinal direction, the radius of the shell was kept fixed while sliding its z-position from the QC to the top of the meristem (Figures S6 and S7). These data gave a general impression of the progressive changes of cellular features in individual or neighboring tissues (depending on bin position) along the z-axis. Again, they confirmed the reduced anisotropy of *bri^TRIPLE^* cells from early on (Figure S6A-C), but also suggested an earlier transition to volumetric growth than in wildtype (Figure S7A). Because of the increased width of *bri^TRIPLE^* roots, especially the wider stele (Figure S8A), this transition was offset in the strictly per-position comparisons (Figure 4A-C). However, when sliding window curves were aligned with the average position of pericycle cells as reference point for the border between the smaller vascular and the considerably bigger ground tissue cells (Figure 4D-F), it became evident that cells in *bri^TRIPLE^* mutants gained in volume earlier than in wildtype (Figure 4D). This coincided with cells approaching wildtype length (Figure 4E) while maintaining an increased width (Figure 4F). In summary, our analyses confirmed that *bri^TRIPLE^* cells across tissues displayed reduced anisotropy already at the outset, and also suggested that their volumetric expansion preceded wildtype.

**Figure 4.**
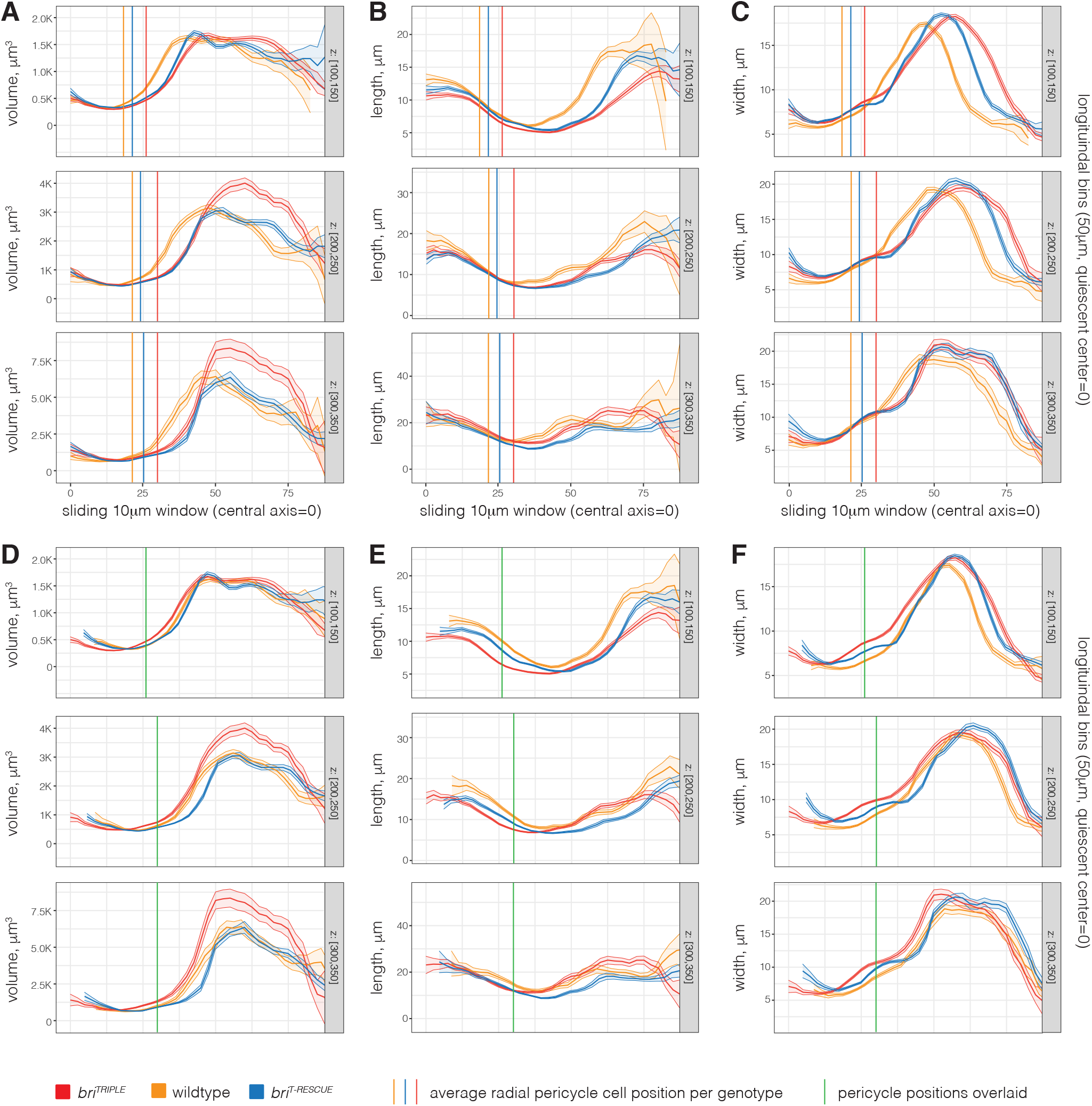
Comparative radial sliding window analysis of standardized root models. (A-C) Extract of quantitative features of each simplified root meristem model (Figures S4 and S5). Models were analyzed using concentric 10 *μ*m thick and 50 *μ*m high cylindrical shells with increasing radius (x-axis) fixed z-position (indicated on the right). Average parameters were calculated by taking into account cells whose centers (after realignment) fell into the corresponding shell. The graphs indicate average cell volume (*μ*m^3^) (A), length (*μ*m) (B) and width (*μ*m) (C) for 11-12 roots per genotype combined. Shaded regions indicate +/− standard error of the mean. Note the displacement of the *bri^TRIPLE^* measurements as compared to wildtype, due to the increased diameter of *bri^TRIPLE^* meristems. Vertical lines indicate the average position of pericycle cells with respect to the central xylem axis in the different genotypes 100*μ*m, 200*μ*m or 300*μ*m above the QC. (D-F) Graphs from (A-C) with curves shifted in order to center them on the average pericycle position (green vertical lines). Note the comparatively premature transition of *bri^TRIPLE^* cells to volumetric growth with growing distance from the QC.

### Brassinosteroid signaling is required for precise cell division orientation

One conspicuous feature of *bri^TRIPLE^* meristems is their disorganized appearance (Figure 2A), which can be mainly attributed to the occurrence of oblique cell division planes that break up the stereotypic, precise alignment of cell files observed in wildtype (Figure S9A-D). For an organ-wide quantification of this phenomenon, we determined the orientation of the triangles that defined the cell surfaces in the 3D meshes of the segmented roots, by evaluating the angle between their normal vector and the z-axis. Plotting the distribution of angles (weighted by triangle area) demonstrated that in wildtype, cell walls are overwhelmingly oriented in anticlinal or periclinal orientation as expected (Figure 5A). By contrast, the distribution of cell wall orientation angles was substantially wider in *bri^TRIPLE^* meristems (Figure 5B and D), matching their visual appearance. A focused analysis on anticlinal cells walls (angles smaller than 45°) corroborated the notion that this deviation is frequent and statistically significant (Figure 5E). Such imperfect alignments also sometimes appeared to cause a “ballooning” phenomenon, where neighboring cells in a file that were separated by an oblique division plane slid next to each other as they elongated (Figure 5F-H). In *bri^TRIPLE^* meristems, such imperfectly aligned cell files could be observed routinely and sometimes created the deceptive impression of the presence of additional cell files. Yet, we also observed late formative divisions in *bri^TRIPLE^* meristems (Figure S2G). These findings suggest that extra cell files in cross sections (Figure S10A) (Kang et al., 2017) could reflect a combination of ballooning effects and late or isolated periclinal divisions. However, the consistently wider stele in *bri^TRIPLE^* meristems (Figure S8A), the steady increase of cell number per z-axis position from the start (Figure S10B), and the increased number of cell files in the mature region (Figure S10C) reiterate the major contribution of supernumerary formative divisions. In summary, the analyses suggest that although cell proliferation capacity is not impaired in *bri^TRIPLE^* meristems, the precision of cell divisions is.

**Figure 5.**
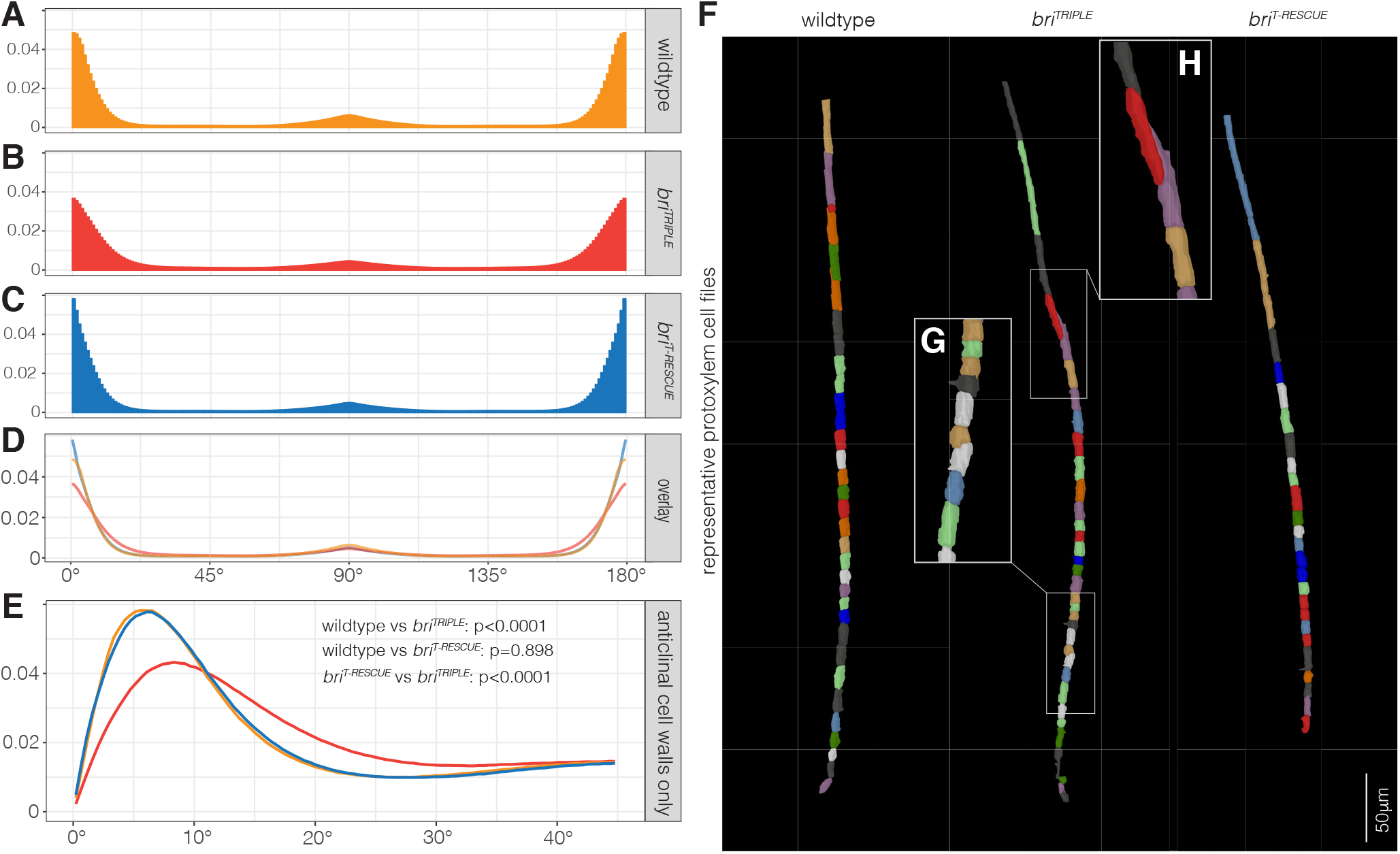
Analysis of global cell wall orientation. (A-C) Distribution of angles between z-axis and vectors normal to the 3D-meshes triangles per unit of solid angle (weighted by triangle area) of 11-12 wildtype (A), *bri^TRIPLE^* (B), and *bri^T-RESCUE^* (C) roots. 90° signifies normal vector perpendicular to the z-axis, 0° signifies normal vector parallel to the z-axis. (D) Superimposition of the three plots highlights the wider distribution of angles in *bri^TRIPLE^* cells, indicating more surfaces that are poorly aligned with the main axes. (E) Distribution of angles (weighted by triangle area) limited to triangles with <45°. The spread of the distribution away from =0 was estimated as the median of the distribution (weighted by triangle area and considering only triangles with theta <45°), which was evaluated for each root separately. Indicated p values were obtained by Welch two-sided T-tests of median grouped by genotype. (F-H) Representative protoxylem cell file segmentations (F), illustrating oblique cell division planes in *bri^TRIPLE^* meristems (G) and the consequent “ballooning effect” of cells sliding next to each other, “out of file “, upon elongation (H).

### Phloem-specific BRI1 imparts an intermediate compensatory state on *bri^TRIPLE^* meristems

Interestingly, despite the macroscopically nearly wildtype growth vigor of *bri^T-RESCUE^* roots (Figure S2A) and the essentially normal morphology of their mature cells (Figure 1A-E), their meristems displayed distinct abnormalities (Figure 2A) that became more apparent in the 3D characterization (Figure 3F, Movie S6). Throughout the analyses, *bri^T-RESCUE^* cells were overall quantitatively situated between *bri^TRIPLE^* and wildtype (Figures 2B-D and S4–S8). However, they generally appeared more similar to *bri^TRIPLE^* in the meristematic region, and transitioned to a more wildtype-like state as cells elongated and differentiated. That is, although cells in *bri^T-RESCUE^* meristems appeared initially less anisotropic than in wildtype, they eventually caught up and did not undergo the comparatively earlier transition to volumetric growth observed in *bri^TRIPLE^* meristems, but rather approached the growth trajectory observed in wildtype (Figures 4D). In line with the overall visual impression of normalized tissue organization as compared to *bri^TRIPLE^* meristems (Figure 2A), cell wall orientations were largely restored with respect to the main axes (Figure 5C-D) and anticlinal cell wall orientations were indistinguishable from wildtype (Figure 5E). Finally, although *bri^T-RESCUE^* meristems also displayed supernumerary periclinal or formative divisions (Figures S8 and S10A-B), this only translated into a modest increase of cell files in the mature roots (Figure S10C). This could be mainly attributed to extra vascular cell files, matching the in-between diameter of the stele (Figure S8A), whereas the phenomenon was less pronounced in the bigger ground tissue cells (Figure S10A and C). In combination with the restored anisotropy (Figure 1B-D), this resulted in a mature root diameter that was close to wildtype (Figure 1A). In summary, the data suggest that restriction of brassinosteroid signaling to the developing phloem cell files predominantly impacts the trajectory of cell growth and differentiation and normalizes the cellular anatomy even in distant brassinosteroid-blind cell files through non-cell-autonomous functions.

### The intermediate compensatory state is associated with a unique transcriptomic signature

In order to understand the role of brassinosteroids in determining cell morphogenesis and discern local differences underlying the non-cell-autonomous rescue, we performed single-cell mRNA sequencing (scRNAseq) of *bri^TRIPLE^* and *bri^T-RESCUE^* meristems. After quality control and filtering, 5,767 and 6,724 high quality cells, respectively, were obtained and next compared to a wildtype root meristem dataset of 5,061 cells produced on the same platform (Wendrich et al., 2020). Unsupervised clustering identified 28 distinct groups of cells, which could be further sorted into 38 subclusters (Figure 6A-C, Table S1-S3) based on known tissue-as well as stage-specific wildtype marker genes (Wendrich et al., 2020). The relative abundance of cells within the clusters was overall comparable to wildtype, although several subclusters contained very few cells (Figure S11A). The transcriptomics data suggest that individual tissues differentiate correctly in the absence of brassinosteroid signaling, and despite their (relatively mild) differentiation defects (Cano-Delgado et al., 2004; Kang et al., 2017), vascular tissues of *bri^TRIPLE^* roots could be readily identified. Globally, the differences between *bri^T-RESCUE^* and *bri^TRIPLE^* were quantitatively less pronounced than between either of those two genotypes and wildtype, corroborating earlier observations from bulk mRNA sequencing (Graeff et al., 2020). Thus, the phenotype of *bri^T-RESCUE^* meristems was associated with a transcriptomic signature that was intermediate between *bri^TRIPLE^* and wildtype.

**Figure 6.**
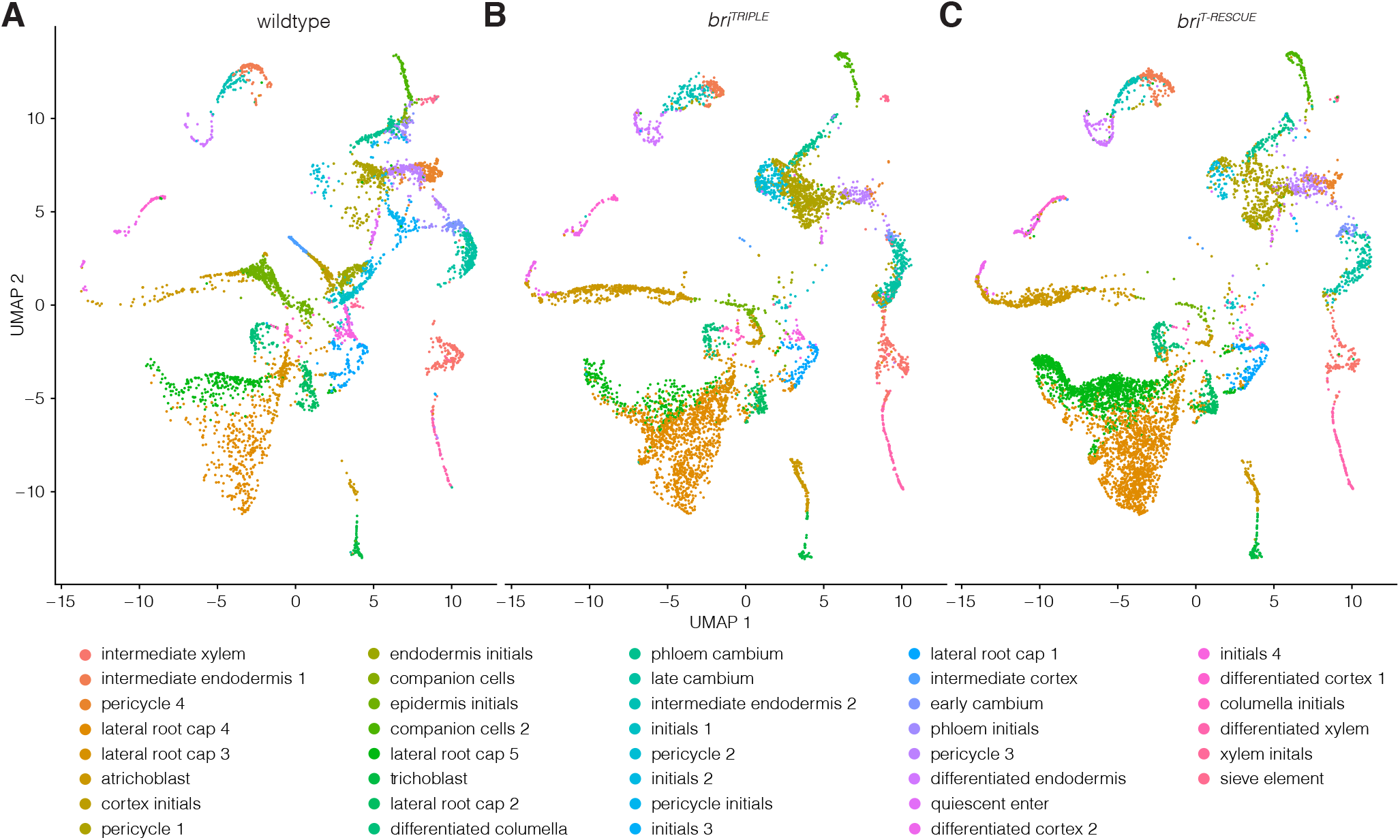
Single cell mRNA (scRNAseq) sequencing overview. (A-C) Uniform Manifold Approximation and Projection (UMAP) of wildtype (A), *bri^TRIPLE^* (B), and *bri^T-RESCUE^* (C) single cell transcriptomes (see Tables S1-S3). Colours indicate the assigned cell identities based on established cell type- and stage-specific marker genes (Wendrich et al., 2020).

To specifically monitor known brassinosteroid-responsive genes, we compared our data to two reference sets. On the one hand, we specifically investigated the overlap with a high confidence list of generically brassinosteroid-regulated genes (Liu et al., 2020); on the other hand, we looked at a set of particularly pertinent genes that were reported to respond to brassinosteroid specifically in root tips (Chaiwanon and Wang, 2015). For a first analysis, the stage-specific scRNAseq subclusters were merged to represent the thirteen cell types (Wendrich et al., 2020) (Figures S11B and S12A-C, Table S4). Considering the reduced sequencing depth of scRNAseq as compared to conventional bulk transcriptomics, the overlap was substantial. For example, of the 1,385 brassinolide-downregulated and 2,312 brassinolide-upregulated genes reported for the root tip of a brassinosteroid biosynthesis mutant (Chaiwanon and Wang, 2015), 576 and 1,218 transcripts, respectively, were detected among the differentially expressed genes in at least one cluster comparison (adj. p<0.05) (Figure S13A-B, Tables S5 and S6). Among them, 525 and 1,040, respectively, showed differential expression between *bri^TRIPLE^* and wildtype (Figure S14A-B). These findings matched the expected behavior of known brassinosteroid-responsive genes in *bri^TRIPLE^* roots, i.e. brassinosteroid-downregulated genes were typically over-expressed in the mutant, whereas brassinosteroid-upregulated genes typically showed reduced expression levels. A substantial portion (~39-43%) of the affected genes were also differentially expressed in *bri^T-RESCUE^* meristems as compared to *bri^TRIPLE^* meristems (Figures S13A-B and S14A-B), and consistently resembled the expression in wildtype. However, an equally large portion (~43-48%) of genes was still differentially expressed between *bri^T-RESCUE^* and wildtype meristems (Figure S13A-B). Because brassinosteroid-responsive genes in our set were particularly prominent in the cortex, we tested whether this skewed our analyses. However, our observations also held up when cortex cells were excluded from the comparison (Figure S13C-D). Finally, the trends described above were confirmed by similar comparisons with the generic set of brassinosteroid-responsive genes (Liu et al., 2020) (Figures S13E-F and S15). In summary, the *bri^T-RESCUE^* genotype again represented an intermediate state between *bri^TRIPLE^* and wildtype.

### Brassinosteroid-responsive genes can be controlled via non-cell-autonomous signals

Including the subset of 732 brassinosteroid-responsive reference genes (Chaiwanon and Wang, 2015), the *bri^TRIPLE^* and *bri^T-RESCUE^* scRNAseq transcriptomes displayed 2,157 genes that were differentially expressed in at least one cell type (adj. p<0.05). In depth comparison of tissue-specific differential gene expression yielded some insight into the nature of the rescue. For instance, we found that 310 genes showed robust differential expression in phloem tissues, however only a minority (n=32) were specific for the phloem and of those even fewer (n=9) were also known brassinosteroid-regulated genes. This pattern was also observed in the subcluster analysis (p<0.01), where 63 of 278 genes that were differentially expressed in sieve elements were also known as brassinosteroid-responsive. Even less were sieve-element-specific (10 and 109, respectively), but included a few known protophloem sieve element markers such as *ALTERED PHLOEM DEVELOPMENT* (Bonke et al., 2003), *BREVIS RADIX* (Rodriguez-Villalon et al., 2014) or *CLAVATA3/EMBRYO SURROUNDING REGION-RELATED 26* (CLE26) (Anne et al., 2018). The differentially expressed phloem and sieve element transcripts spanned a variety of encoded proteins, which might include factors that are involved in the generation or transmission of the non-cell-autonomous signal and could thereby aid in its discovery.

Importantly, in *bri^T-RESCUE^* meristems, the expression of many brassinosteroid-responsive genes was not only normalized in the phloem, but also in directly neighboring as well as distant tissue layers (Figures S14 and S15). This was also evident in the more fine-grained analysis across the stage-specific clusters (Figure S16). That is, the expression of brassinosteroid-responsive genes was not only fully normalized in sieve elements and phloem initials as expected, but also other clusters like xylem initials or intermediate cortex. Moreover, in clusters in which a higher number of brassinosteroid-responsive genes could be detected, the expression of a sizeable portion was normalized to the same degree as in the overall analyses (Figure S16A-B). These results suggest that brassinosteroid can control many genes indirectly, through non-cell-autonomous signals.

### Arabinogalactan-encoding genes are sentinels of brassinosteroid-dependent cellular anisotropy

Apart from known brassinosteroid-responsive genes, transcripts whose expression was fully or partially normalized in *bri^T-RESCUE^* meristems can be considered a systemic readout of such non-cell-autonomous signals. We sought to determine whether they comprise particular gene classes that are differentially expressed throughout tissues in a coherent manner. However, gene ontology analyses did not reveal any consistent pattern. Yet, genes encoding arabinogalactan proteins and peptides (AGPs) clearly stood out because of their robust, comparatively high expression in *bri^T-RESCUE^* tissues as compared to the *bri^TRIPLE^* mutant (Figure S17A, Table S7). This observation is in line with the proposed role of AGPs in cell growth in various systems (Seifert and Roberts, 2007; van Hengel and Roberts, 2002; Willats and Knox, 1996), and the positive correlation between their expression level and cell elongation (Pacheco-Villalobos et al., 2016). By contrast, differential expression of the classic cell expansion genes, the expansins, which facilitate cell growth by promoting cell wall loosening (Cosgrove, 2005), was only observed sporadically (Figure S17B, Table S8). Moreover, those expansins that displayed differential expression were nearly exclusively over-expressed in the *bri^TRIPLE^* meristems, which might reflect the premature transition to volumetric growth described above. These observations were corroborated in the subcluster analysis, where most *AGPs* (23/32) showed differential expression between *bri^TRIPLE^* mutant and wildtype or *bri^T-RESCUE^* tissues, and generally also displayed stronger expression at later stages of tissue ontogeny (Figure 7A). This included many *AGPs* (15/23) that were part of the brassinosteroid-responsive set (Chaiwanon and Wang, 2015). Again, such strong and spatially graded expression differences were not observed for expansins (Figure 7B), although several (10/35) have also been described as brassinosteroid target genes. In light of our finding that volumetric growth is not affected in *bri^TRIPLE^* mutants, but cellular growth orientation is, *AGPs* could therefore be considered sentinels of brassinosteroid-dependent cellular anisotropy and represent the most prominent read-out of non-cell-autonomous brassinosteroid functions.

**Figure 7.**
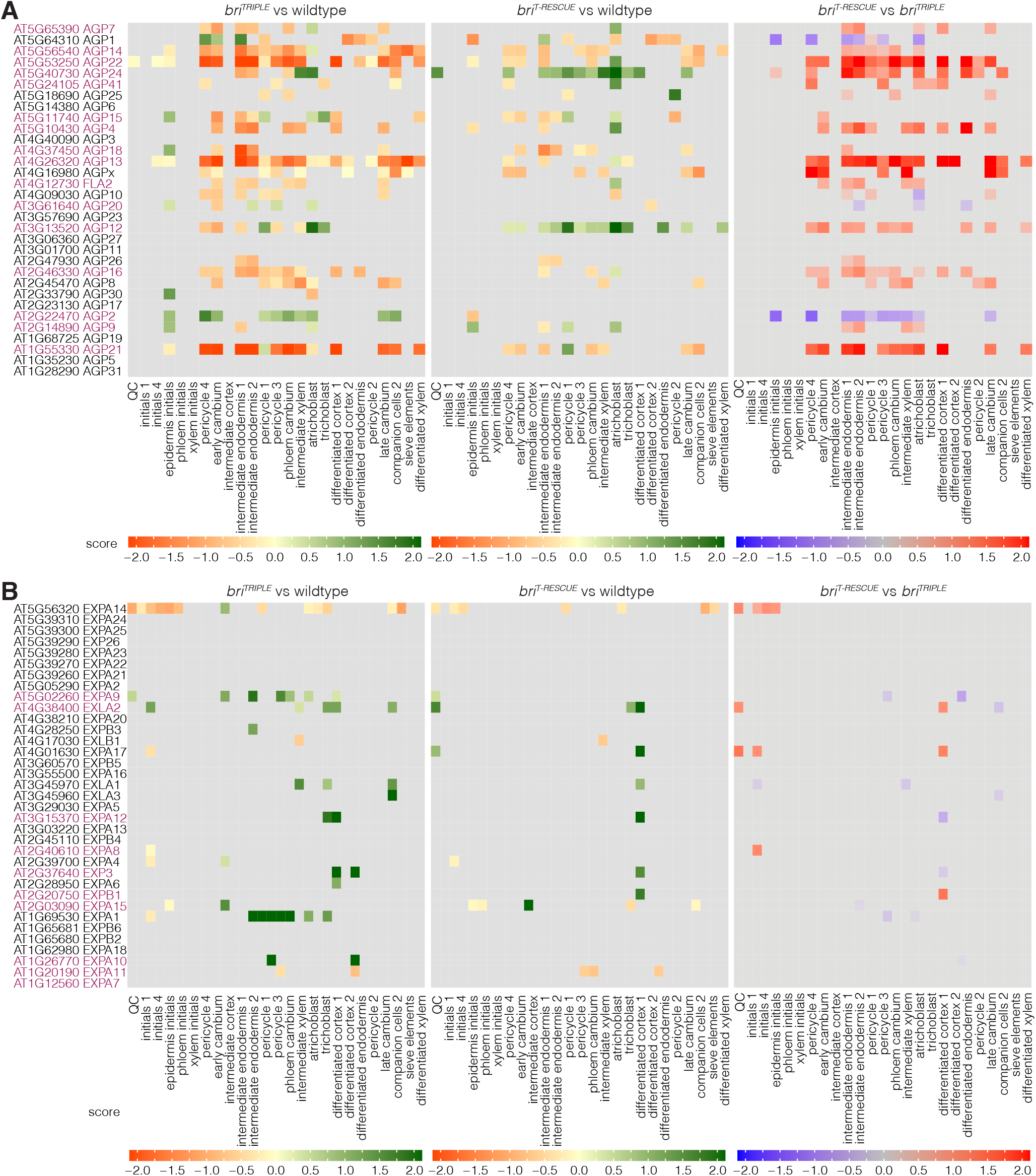
*AGP* gene expression is a sentinel of cellular anisotropy. (A-B) Heatmap indicating the differential gene expression score of AGPs (A) or expansins (B) in scRNA-seq subclusters and ordered with respect to stage-specific markers in increasing distance from the QC. Known brassinosteroid-responsive gene family members (Chaiwanon and Wang, 2015) are highlighted in purple.

## DISCUSSION

### Brassinosteroid signaling controls cellular growth orientation

In summary, our combined analyses reveal that brassinosteroid signaling in root development is essential for the precise orientation of cell division planes and the extent and timing of anisotropic cell expansion. Our conclusions highlight the strength of quantitative 3D digital single-cell analyses and show that they are necessary to arrive at non-obvious and quantitatively confirmed conclusions.

The observed absence of effects on cell volume and proliferation capacity versus the statistically significant effects on cellular anisotropy and division plane orientation open new perspectives on previous analyses. For instance, the notion that cell division frequency is reduced in roots of brassinosteroid signaling mutants was largely deduced from the abundance of cell division markers relative to 1D meristem size expressed as the number of meristematic cells in individual cell files. However, this method is prone to error especially when absolute differences are small (Perilli and Sabatini, 2010), which might explain why effects on meristem size are not consistent across studies (Chaiwanon and Wang, 2015; Gonzalez-Garcia et al., 2011; Hacham et al., 2011; Kang et al., 2017). Despite the normal frequency of dividing cells, our estimations suggest that the number of cells produced per time unit is indeed reduced in *bri^TRIPLE^* roots. This might reflect slowed down root growth due to the production of extra cell files (De Rybel et al., 2013) as previously proposed (Kang et al., 2017) or slower progression of the cell cycle (Gonzalez-Garcia et al., 2011; Hacham et al., 2011), but it could also be a consequence of sub-optimal cell-cell communication due to the disorganized meristem. High resolution live imaging of *bri^TRIPLE^* roots that express pertinent dynamic cell cycle markers might be required to gain ultimate insight. Furthermore, based on 1D cortex cell length measurements and modeling, it has been proposed that brassinosteroid signaling is required to set absolute cell size (Pavelescu et al., 2018). Our 3D data are consistent with this proposition with regard to cell elongation, but also show that the sizer mechanism applies to cellular anisotropy rather than a reduction of cell size in terms of volume. The same study suggested that brassinosteroid signaling suppresses a timer mechanism of cell elongation (Pavelescu et al., 2018), which resonates with the premature onset of volumetric growth in *bri^TRIPLE^* meristems observed here. Interestingly, this result matches the graded activity of canonical brassinosteroid signaling in the root meristem, which increases as cells start to elongate and differentiate (Chaiwanon and Wang, 2015). Thus, timing the transition to anisotropic growth might be a primary effect of brassinosteroid signaling in the root, and also an effect that can be conveyed by brassinosteroid-dependent non-cell-autonomous signals.

### Brassinosteroid-controlled genes can respond to non-cell-autonomous brassinosteroid action

The latter conclusion is derived from our finding that the systemic rescue of *bri^TRIPLE^* roots through phloem-specific brassinosteroid signaling manifests in a normalized anatomy of mature cells. Yet, it is also associated with an intermediate meristem phenotype. Although *bri^T-RESCUE^* meristems are thus not entirely resembling wildtype meristems, the cell-aligning effects of brassinosteroid signaling specifically in the developing phloem can obviously normalize the anatomy of both adjacent and distant brassinosteroid-blind cells to a degree that is sufficient to impart nearly wildtype root growth vigor. In line with this key finding, the single-cell mRNA sequencing data demonstrate that many brassinosteroid-controlled genes can respond to the postulated non-cell-autonomous signals. In this context, the expression of genes encoding AGPs was a particularly robust systemic readout, and our data therefore qualify *AGPs* as sentinels of brassinosteroid-dependent cellular anisotropy rather than of cell growth *per se*. *AGPs* can therefore be of use to trace the activity of non-cell-autonomous brassinosteroid-triggered signals. Given the strong effect of the rescue, one could assume that these signals are self-reinforcing but need to be activated by brassinosteroid perception in the first place. Such propagation could explain the far reach of the signal, including its apparently growing impact as cells transition to elongation and differentiation across cell layers although the developing phloem sieve elements differentiate very early and no longer express the brassinosteroid receptor once they differentiate (Graeff et al., 2020). The present comprehensive analysis of brassinosteroid effects in the root might aid in the discovery of the non-cell-autonomous signals and can also serve as a blueprint for comparative analyses to determine whether brassinosteroid action in the shoot is based on similar principles.

## METHODS

### Plant materials and growth conditions

Seeds were surface sterilized using 3% sodium hypochlorite, sown onto half strength Murashige & Skoog agar medium (0.9% agarose) supplemented with 0.3% sucrose, and stratified for 3 days at 4°C. Plants were grown under continuous white light (intensity ~120 μE) at 22°C. All mutants and marker lines were in the *Arabidopsis thaliana* Columbia-0 (Col-0) wildtype background. The *bri^TRIPLE^* mutant and *bri^T-RESCUE^* lines have been described previously (Graeff et al., 2020).

### Tissue fixation and clearing

For confocal microscopy, roots of 7-day-old seedlings were fixed in 4% PFA in PBS and transferred into a vacuum of 25 to 30 mmHg/Torr for 15 to 30 min, then washed 3 times in PBS for 5 min. For clearing of the samples, seedlings were transferred into *ClearSee* solution (Kurihara et al., 2015; Ursache et al., 2018) for 7 days at 4°C with regular changes of the clearing solution. Cleared roots were stained with 0.25 mg/mL Calcofluor White (CCFW - Sigma, Product No. F3543), washed with *ClearSee* solution and mounted on microscope slides with distancers in *ClearSee* solution that contained 0.3% agarose to prevent moving of the samples.

### Confocal microscopy

Samples were imaged on a Zeiss LSM880 confocal microscope with a 40x objective, using a 405 nm laser for CCFW excitation and recording of the cell wall signal in the 450 to 480 nm range. Tile scans and z-stacks were set to image 500 μm of root meristem or 1000 μm of mature root (located ~1 cm shootward from the root tip), respectively. The distance between the z-scans was set to 0.42 μm for the root meristems and 0.5 μm for the mature roots, with a pin hole diameter of 33 μm. 512 × 512 pixel 16 bit images were recorded with a scanning speed of 2.06 μs per pixel.

### *PlantSeg-MorphoGraphX-3DCellAtlas* workflow

*Segmentation of cells in the images:* For segmentation of root micrographs, the channel of the stained cell walls was extracted as 16 bit .tiff image. The *PlantSeg* workflow (Wolny et al., 2020) was used to predict cell boundaries and label the cells in the image stacks. The re-scaling factor for the images was calculated based on their resolution and the model used for cell boundary prediction ([1.70, 2.8, 2.8] to fit root meristems to the ≪confocal_unet_bce_dice_ds2x≫ model; and [1.74, 2.04, 2.04] to fit the mature root images to the ≪confocal_unet_bce_dice_ds3x≫ model). GPU-based prediction was used for cell boundary prediction, followed by segmentation using the GASP segmentation algorithm with beta: 0.5, 2D watershed and a watershed threshold of 0.5. Segments smaller than 1000 were discarded. After segmentation, images were re-scaled with the appropriate factors and saved as .tiff image files. *Creation of 3D meshes from the segmented images and extraction of cell properties*: Segmentations were imported into *MorphoGraphX* software (Barbier de Reuille et al., 2015). 3D meshes of each root were created using the *marching cubes* 3D workflow with a cubesize of 2 and a minimal cell size of 500 voxels. Cell properties were extracted using the *3DCellAtlas* plugin as described (Montenegro-Johnson et al., 2015). Cell types were assigned based on the radial and circumferential coordinate of the cells and manually corrected. Attributes and properties of all cells were exported for comparative analysis. Images of the labelled root meshes using horizontal or transversal cutting planes and of the original image were generated with *MorphoGraphX*. Videos of 3D image stacks and renderings were generated using *Imaris* software.

### Digital single cell analysis using *3DCellAtlas*

Cell types and 3D cell geometries in roots were determined using *3DCellAtlas* (Montenegro-Johnson et al., 2015) implemented within *MorphoGraphX* (Barbier de Reuille et al., 2015) using the methodology described previously (Stamm et al., 2017). Cell type annotation was manually verified and corrected in instances where errors were present. Data describing cell size and shape were exported as CSV files for further statistical analyses.

### Root comparison based on *MorphoGraphX-3DCellAtlas* output

The comparison of cell properties for different genotypes was performed in *R* (4.0.4). Attribute data of the mature root segments were imported and cells with length, width or circumference ≤ 0 μm or a volume < 1000 μm^3^ were removed, leaving a dataset with 9,571 cells. Cell properties were analyzed by the assigned cell type. For statistical comparison, mean values for each cell type were calculated per root and one-sided ANOVA was performed to analyze the significance of the observed variance between roots. Tuckey-HSD tests were performed to compare the variances for each tissue between the genotypes. For comparative analyses of the root meristems, cell types of four roots per genotype were assigned using the *3DCellAtlas* plugin and manually corrected. To compare dimensional properties, each root was re-oriented so that the y axis ran along the longitudinal axis of the root and x- and z-axis defined the horizontal dimensions. The distance of each cell centroid to the center of the SCN was calculated. For cell length, width, circumference, volume and wall area the data distribution was analyzed and cells below the 2.5% quantile and above the 97.5% quantile were discarded to remove segmentation artefacts. Roots were sliced into 5 μm bins along the longitudinal axis, and each cell was assigned to the bin containing its centroid. The dataset contained 46,559 cells in total (13,778 from wildtype, 15,508 from the *bri^TRIPLE^* mutant, and 17,273 from the *bri^T-RESCUE^* line). Cell properties were plotted as scatterplots, a regression curve was fitted to the data for each genotype using a general additive model with a shrinking cubic regression spline interpolation.

### Straightening of 3D meshes, creation of simplified root models and per-position comparisons

The procedure for the straightening and alignment of 3D meshes to produce simplified root meristem models for per-position cell parameter comparisons is described in detail in the separate Supplemental Methods S1 file.

### *10X Genomics* sample preparation, library construction and sequencing

Single-cell sequencing was performed on protoplasts from *bri^TRIPLE^* and *bri^T-RESCUE^* primary seedling roots as described (Wendrich et al., 2020). Briefly, 6-day-old *bri^TRIPLE^* and *bri^T-RESCUE^* roots were cut and protoplasts were isolated, stained for live/dead using 4′,6-diamidino-2-phenylindole (DAPI) at 14 μM final concentration, and sorted on a *BD Aria II* instrument. For each genotype, DAPI-negative protoplasts were selected for further analysis. Sorted cells were centrifuged at 400g at 4°C and resuspended in protoplast solution A to yield an estimated concentration of 1,000 cells/μl. Cellular suspensions were loaded on a *Chromium Single Cell 3’ GEM, Library & Gel Bead Kit* (V3 chemistry, *10X Genomics*) according to the manufacturer’s instructions. Libraries were sequenced on an *Illumina NovaSeq600* instrument following recommendations of *10X Genomics* at the *VIB Nucleomics Core* facility (VIB, Leuven).

### Raw data processing and generation of the gene expression matrix

Demultiplexing of the raw sequencing data was performed with the *10x CellRanger* (version 3.1.0) software *‘cellranger mkfastq’*. The fastq files obtained after demultiplexing were used as the input for *‘cellranger count’*, which aligns the reads to the *Arabidopsis thaliana* reference genome (Ensemble TAIR10.40) using STAR and collapses them to unique molecular identifier (UMI) counts. The result is a large digital expression matrix with cell barcodes as rows and gene identities as columns. Initial filtering in *CellRanger* recovered 10,574 cells for the *bri^TRIPLE^* sample, and 12,335 cells for the *bri^T-RESCUE^* sample. To ensure only high-quality cells were further analyzed, the filtered data provided by *CellRanger* were used. This corresponded to 54,593 mean reads per cell in the *bri^TRIPLE^* sample, and 40,520 mean reads per cell in the *bri^T-RESCUE^* sample.

### Data Analysis (Clustering, Identity Assignment, DEG, and Quality Control)

All analyses were performed in *R* (version 3.6.0). Data pre-processing was performed by the *scater* (version 1.10.1) package following a recommended workflow (Lun et al., 2016). Outlier cells were identified based on two metrics (library size and number of expressed genes) and tagged as outliers if they were five median absolute deviation (MADs) away from the median value of these metrics across all cells. Normalizing the raw counts, detecting highly variable genes, finding clusters and creating tSNE plots was done using the *Seurat* pipeline (version 3.2.3). Differential expression analysis for marker gene identification per subpopulation was based on the non-parametric Wilcoxon rank sum test implemented within the *Seurat* pipeline. Clusters with the same cell annotation based on gene expression analysis were combined to generate a more comprehensible dataset. Samples were merged with the published wildtype root dataset (Wendrich et al., 2020) and annotations were transferred using the *LabelTransfer*-method from the *Seurat* package.

### Visualization of scRNAseq expression data in heat maps

Lists of AGI identifiers for brassinosteroid-responsive genes in the root tip or the whole seedling were obtained from the literature (Chaiwanon and Wang, 2015; Liu et al., 2020). Each gene was referenced to the cluster datafiles and LogFC, adjusted p-value and score were extracted to a new list. If no entry could be found for the respective cluster, N/A values were added. Summarized data were imported into *R* and tile plots for each comparison were created with tile color indicating the score value using the *ggplot2* and *scales* libraries. Gene entries with an adjusted p-value > 0.05 or N/A were displayed in grey. The scales were fixed to a range of - 2 to 2 for better comparability. Bigger datasets were restricted to genes with a significant logFC in at least one cluster. Venn diagrams comparing the overlap between the different datasets were created using the *ggVennDiagram* library.

## SUPPLEMENTAL INFORMATION

**Supplemental Figures S1–S17**

**Supplemental Methods S1:** Creation of standardized, simplified root models.

**Table S1:** Differentially expressed genes in the scRNAseq data in stage-specific subclusters, *bri^TRIPLE^* versus wildtype comparison.

**Table S2:** Differentially expressed genes in the scRNAseq data in stage-specific subclusters, *bri^T-RESCUE^* versus wildtype comparison.

**Table S3:** Differentially expressed genes in the scRNAseq data in stage-specific subclusters, *bri^T-RESCUE^* versus *bri^TRIPLE^* comparison.

**Table S4:** Differentially expressed genes in the scRNAseq data in cell type-specific clusters.

**Table S5:** Comparison of the scRNAseq data with negatively brassinosteroid-responsive genes in the root.

**Table S6:** Comparison of the scRNAseq data with positively brassinosteroid-responsive genes in the root.

**Table S7:** Differential expression of arabinogalactans in scRNAseq clusters.

**Table S8:** Differential expression of expansins in scRNAseq clusters.

**Movie S1:** 3D reconstruction of a mature wildtype root segment.

**Movie S2:** 3D reconstruction of a mature *bri^TRIPLE^* root segment.

**Movie S1:** 3D reconstruction of a mature *bri^T-RESCUE^* root segment.

**Movie S1:** 3D reconstruction of a mature wildtype root meristem.

**Movie S1:** 3D reconstruction of a mature *bri^TRIPLE^* root meristem.

**Movie S1:** 3D reconstruction of a mature *bri^T-RESCUE^* root meristem.

## ACKNOWLEDGEMENTS

We would like to thank Prof. Thomas Berleth for helpful comments on the manuscript, Prof. Gerd Juergens and Dr. Ulrike Mayer for a gift of anti-KN antibody, and Emma Picot for technical support in image analysis. This work was funded by core funding from the University of Lausanne, the Swiss National Science Foundation (grant 310030B_185379, awarded to C.S.H.), The Research Foundation - Flanders (FWO; post-doc fellowship 1215820N, awarded to T.E.), the European Research Council (ERC Starting Grant TORPEDO; 714055, awarded to B.D.R.), the BBSRC (grant BB/S002804/1 to G.W.B.), and the Deutsche Forschungsgemeinschaft (DFG; post-doctoral fellowship GR 5009/1-1, awarded to M.G.).

---

**Figure S1.**
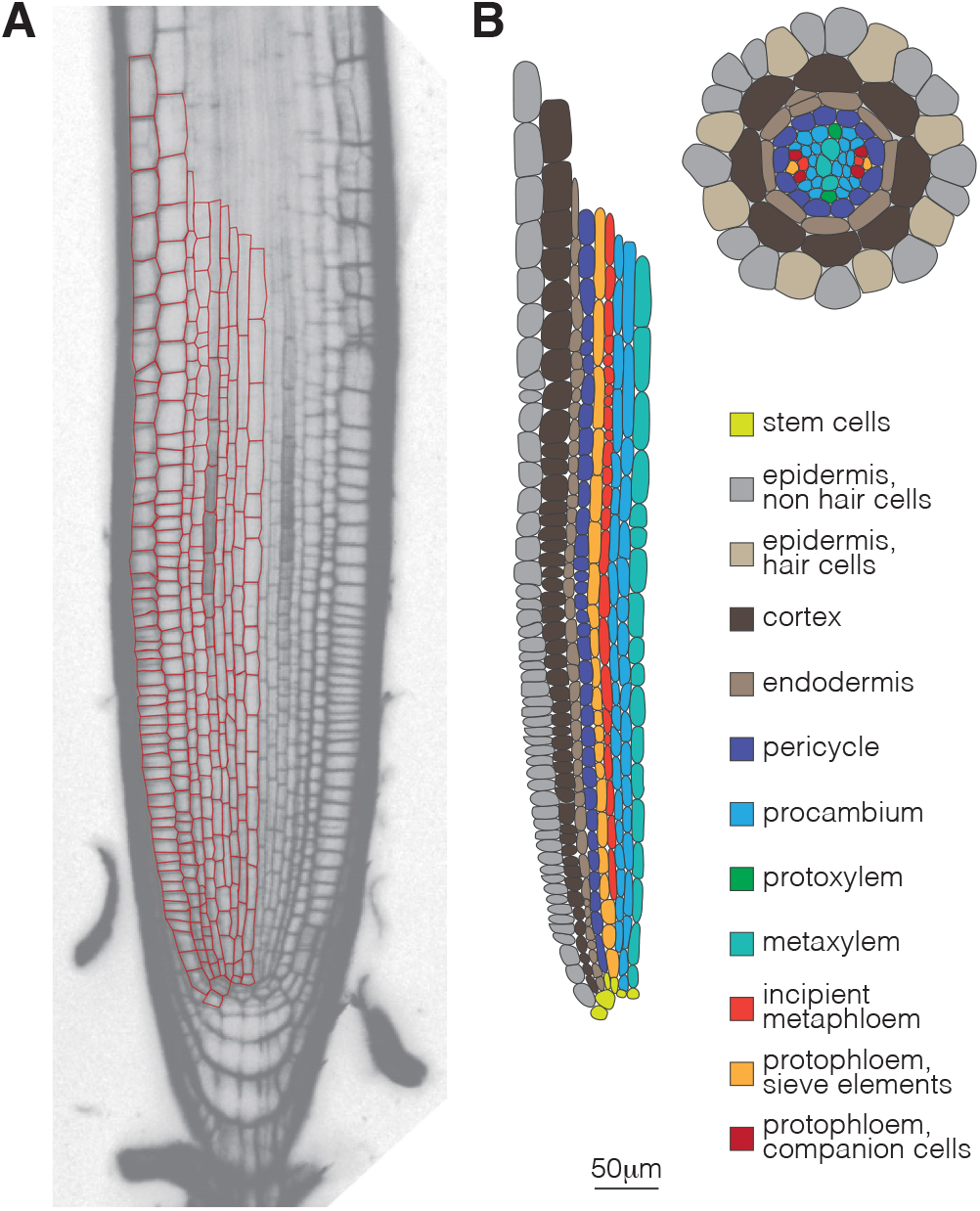
Schematic presentation of Arabidopsis root meristem organization. (A) Transversal confocal section through the center of a representative 7-day-old Arabidopsis root meristem, oriented from phloem pole to phloem pole, merged with cell outlines (red) for one half. (B) Schematic presentation of the tissues in the root shown in (A), labeled for cell types, and a corresponding schematic of their radial arrangement.

**Figure S2.**
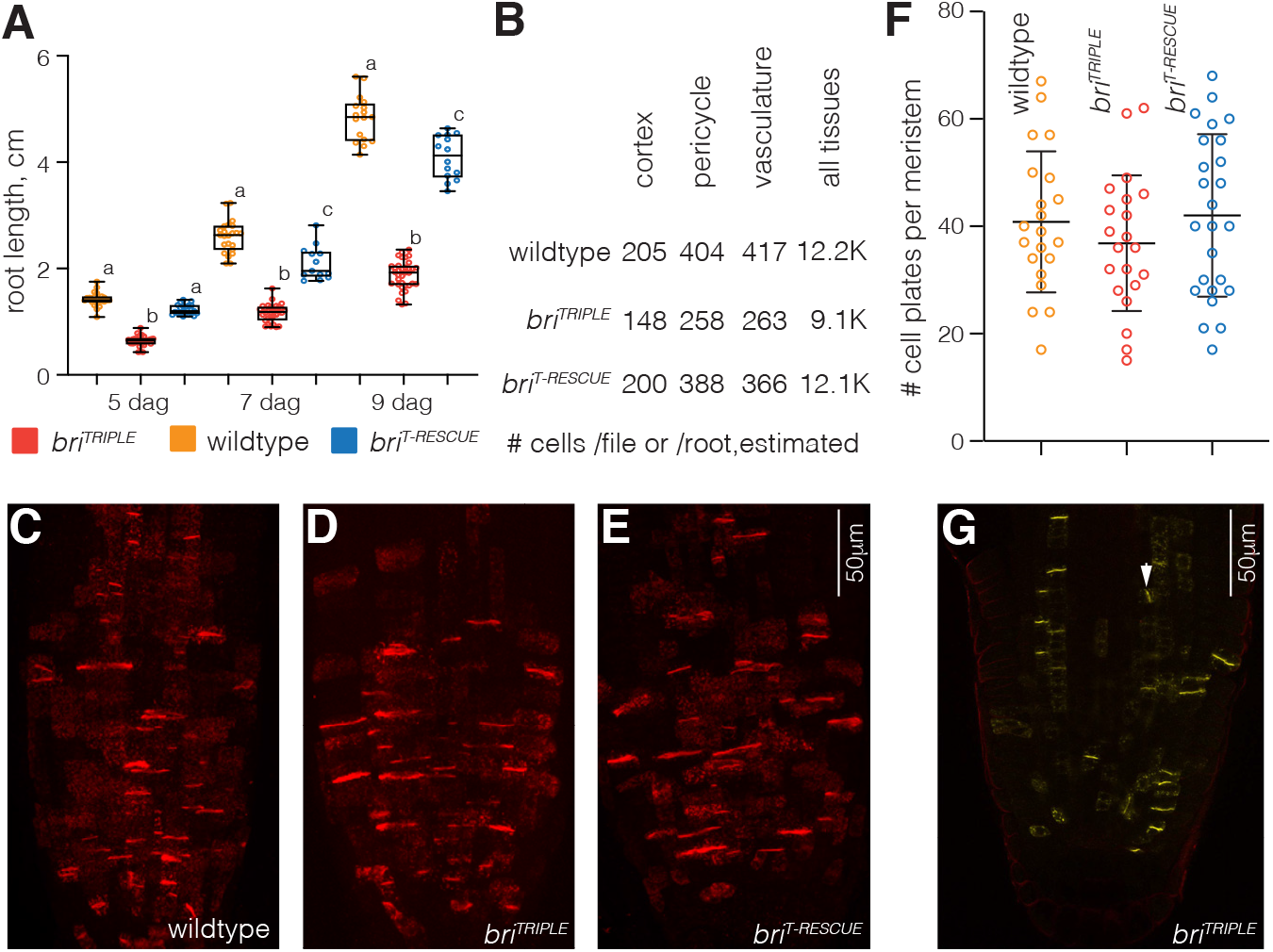
Quantification of cell proliferation. (A) Primary root length of indicated genotypes at 5, 7 and 9 days after germination (dag). Box plots display 2nd and 3rd quartiles and the median, bars indicate maximum and minimum. Statistical significance was determined by ordinary one-way ANOVA. Statistically significant different groups are indicated by different lowercase letters. (B) Estimated number of cells per cell file of indicated tissue or total root mature root cell number, calculated from the average root length (A) and the average cell lengths obtained for mature cells or average cell number per mature segment (Figure 1B). (C-E) Confocal microscopy images of root meristems from indicated genotypes, stained with anti-KN antibody (red fluorescence), showing the maximum projection of the 3D stacks from an entire meristem each. Cell plates of dividing cells are recognizable as sharp KN signal accumulations. (F) Dividing cells per meristem as deduced from cell plates detected by anti-KN immunostaining. Bars indicate the mean and standard deviation. Differences between genotypes were not significant (one-way ANOVA). (G) As in D, for a single slice from a 3D stack. The arrowhead points out a late periclinal division.

**Figure S3.**
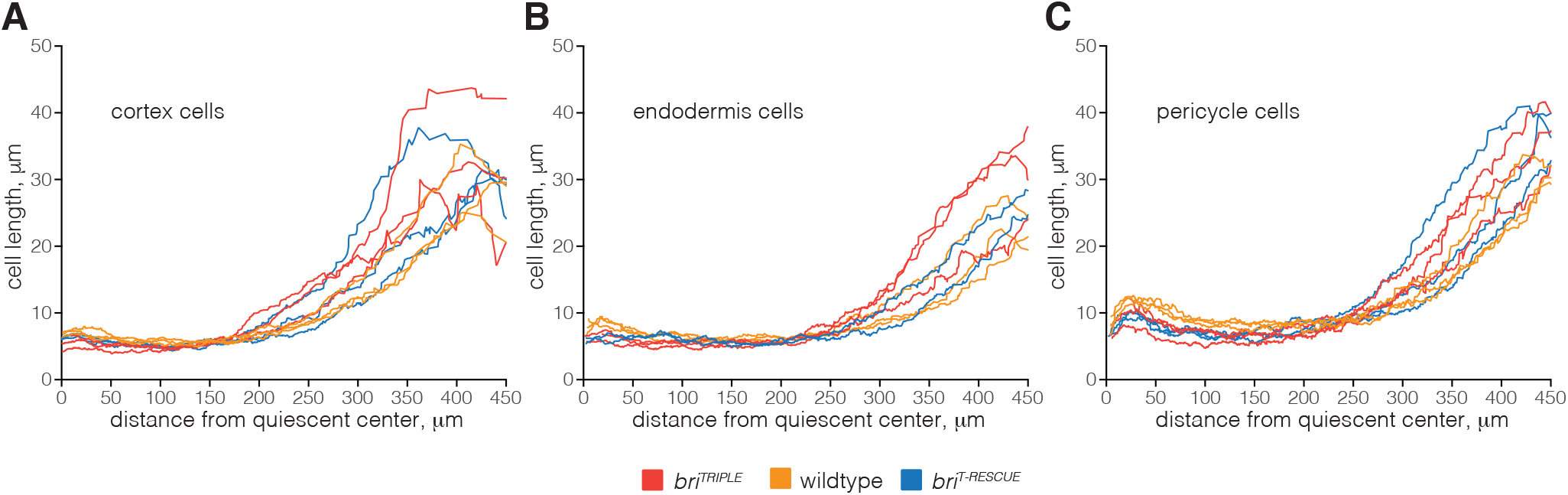
Root growth variability. (A-C) Cell length as a function of distance from the QC, for cortex (A), endodermis (B) and pericycle (C) cell files for a few selected individual roots, obtained through the *PlantSeg-MorphoGraphX-3DCellAtlas* pipeline. Note the substantial variability within and between genotypes.

**Figure S4.**
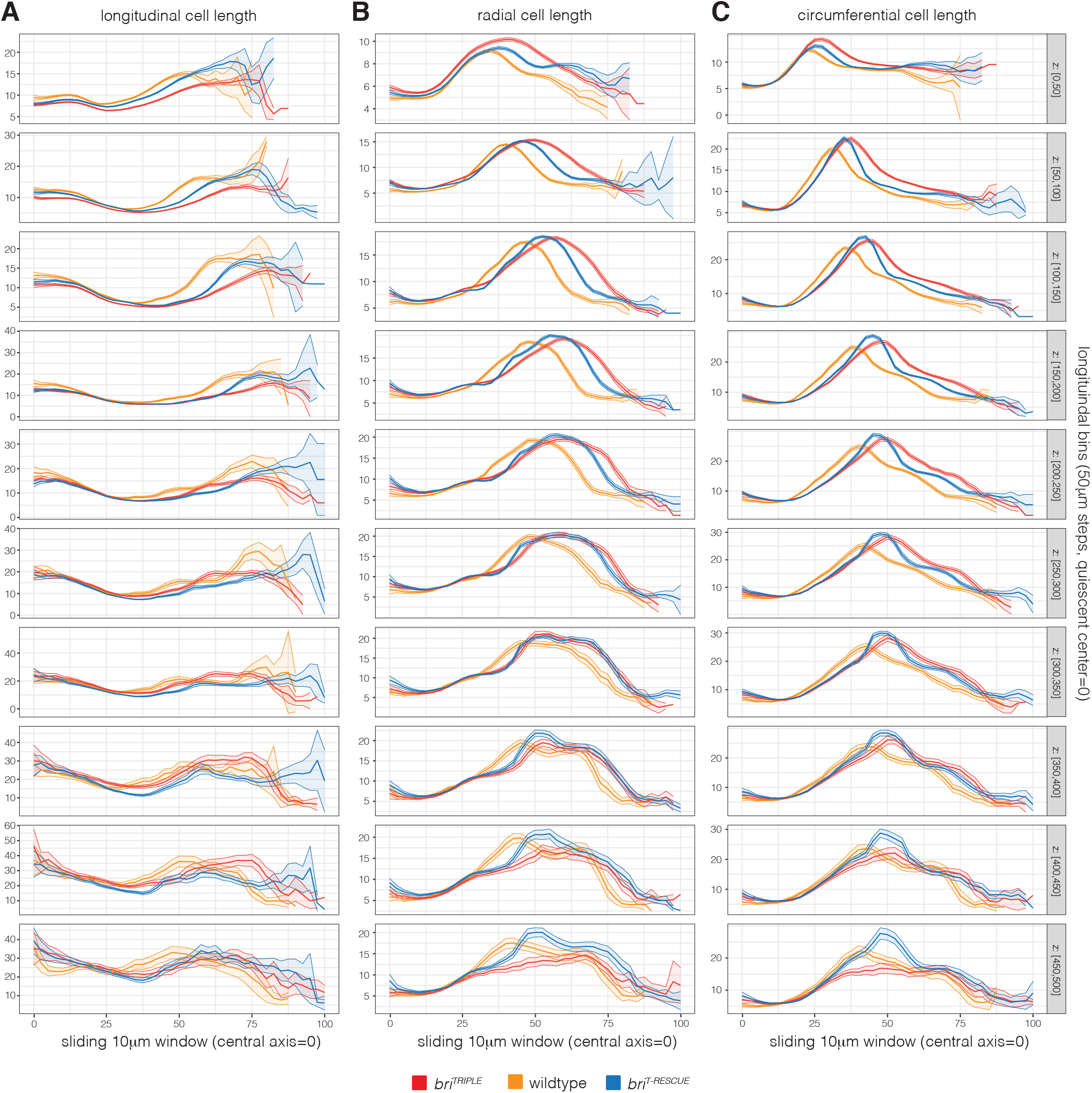
Comparative radial sliding window analysis of standardized root models 1. Quantitative features of each simplified root meristem model using concentric 10 *μ*m thick and 50 *μ*m high cylindrical shells with increasing radius (x-axis) fixed z-position (indicated on the right). Average parameters were calculated by taking into account cells whose centers fell into the corresponding shell. The graphs indicate average longitudinal cell length (*μ*m) (A), cell width in the radial dimension (*μ*m) (B) and cell width in the circumferential dimension (*μ*m) (C) for 11-12 roots per genotype combined. The shaded regions indicate +/− standard error of the mean. Note the displacement of the *bri^TRIPLE^* measurements as compared to wildtype on the x axes, due to the increased diameter of *bri^TRIPLE^* meristems.

**Figure S5.**
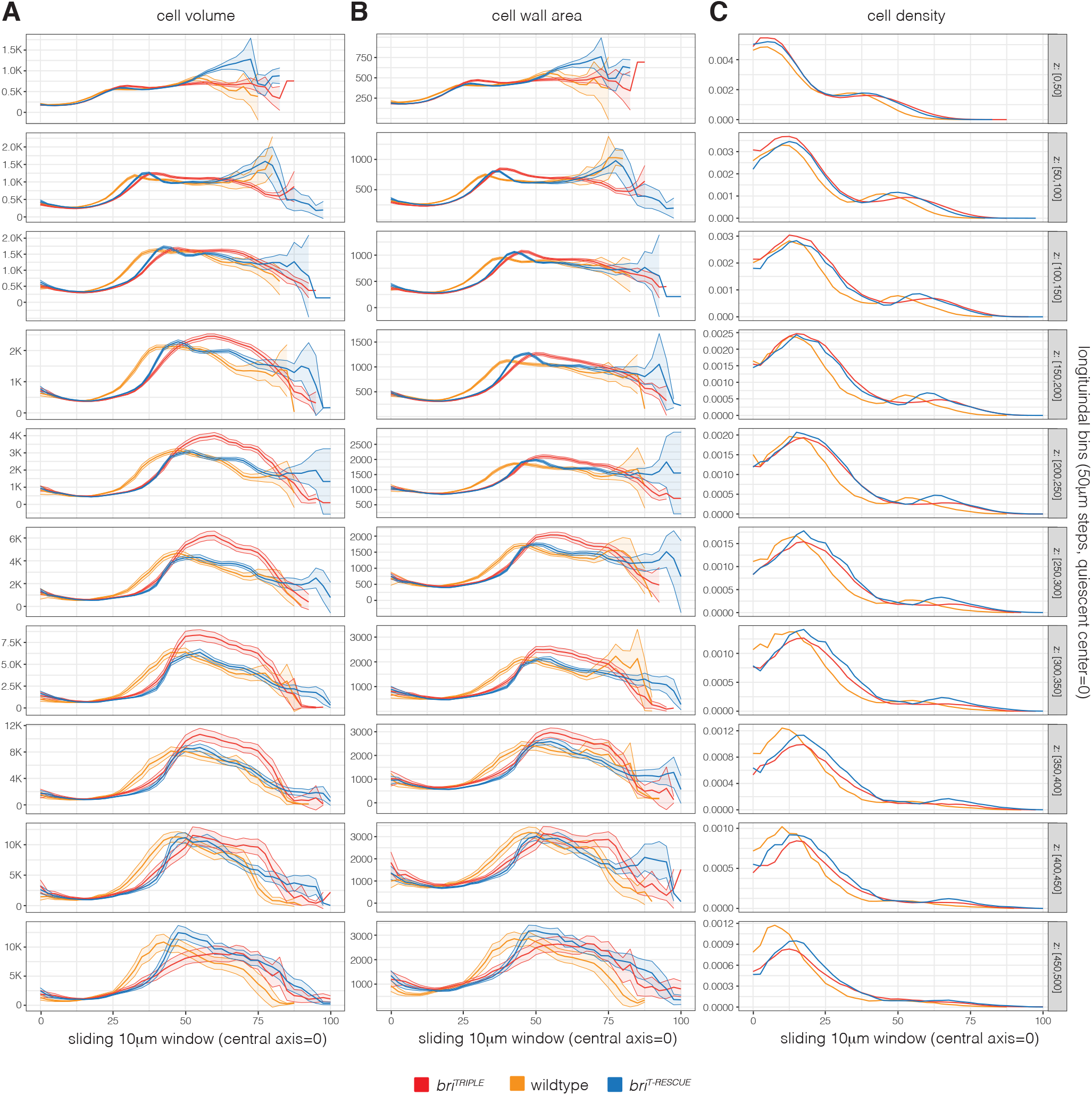
Comparative radial sliding window analysis of standardized root models 2. Continuation of the analysis presented in Figure S4, indicating cell volume (*μ*m^3^) (A), cell wall area (*μ*m^2^) (B), and cell density (cells/*μ*m^3^) (C) for 11-12 roots per genotype combined. The +/− standard error of the mean is indicated where applicable. Note the displacement of the *bri^TRIPLE^* measurements as compared to wildtype on the x axes, due to the increased diameter of *bri^TRIPLE^* meristems.

**Figure S6.**
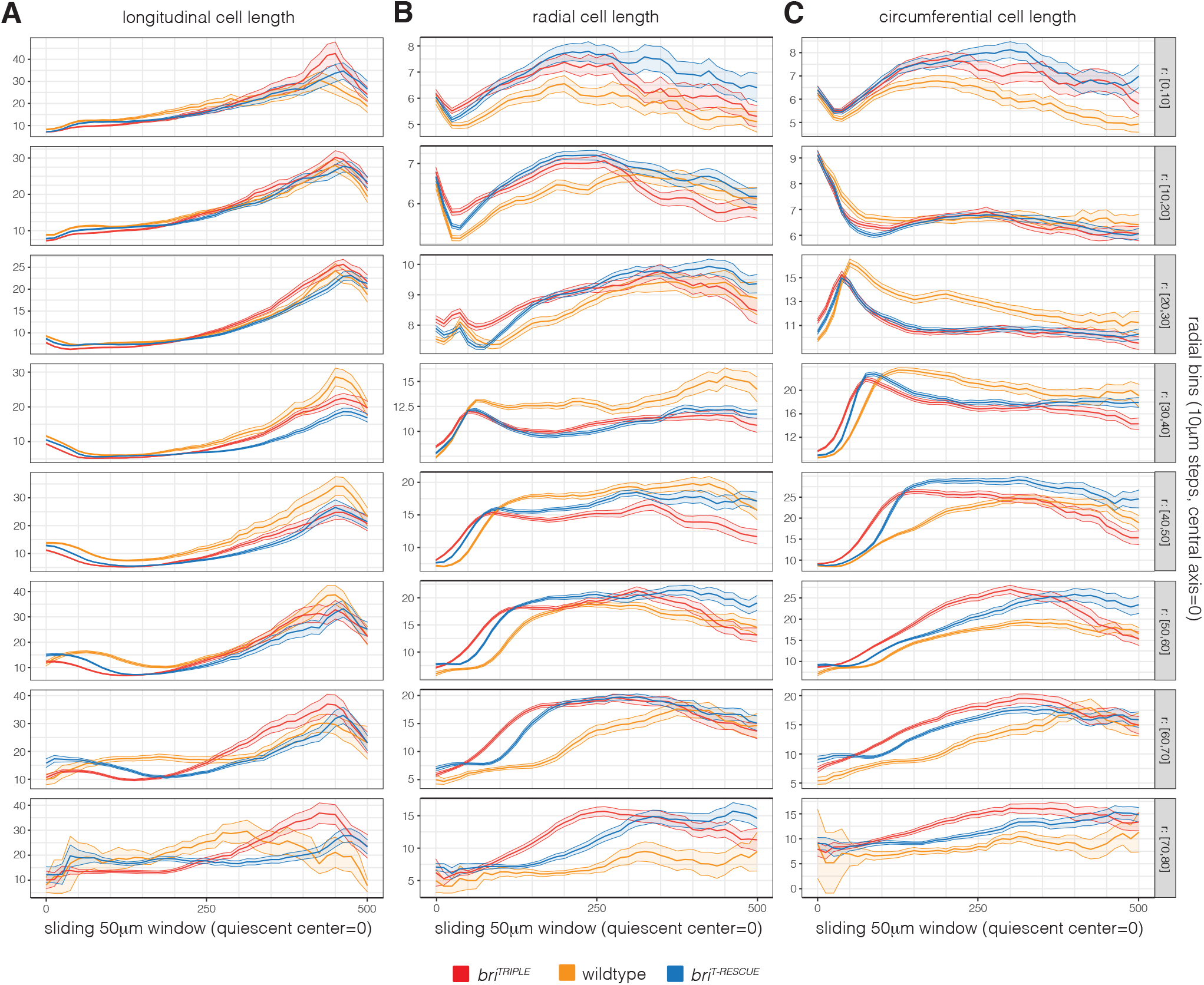
Comparative longitudinal sliding window analysis of standardized root models 1. Sliding window analysis similar to Figure S4, for a 10 *μ*m thick and 50 *μ*m high cylindrical shell (centered on the z-axis) with fixed radius (indicated on the right) and z-position sliding from the QC to the elongation-differentiation zone (x-axis). The graphs indicate average longitudinal cell length (*μ*m) (A), cell width in the radial dimension (*μ*m) (B) and cell width in the circumferential dimension (*μ*m) (C) for 11-12 roots per genotype combined. Shaded regions indicate the +/− standard error of the mean. Note that curves within single panels are not always representing similar tissues, because of the wider stele of *bri^TRIPLE^* meristems.

**Figure S7.**
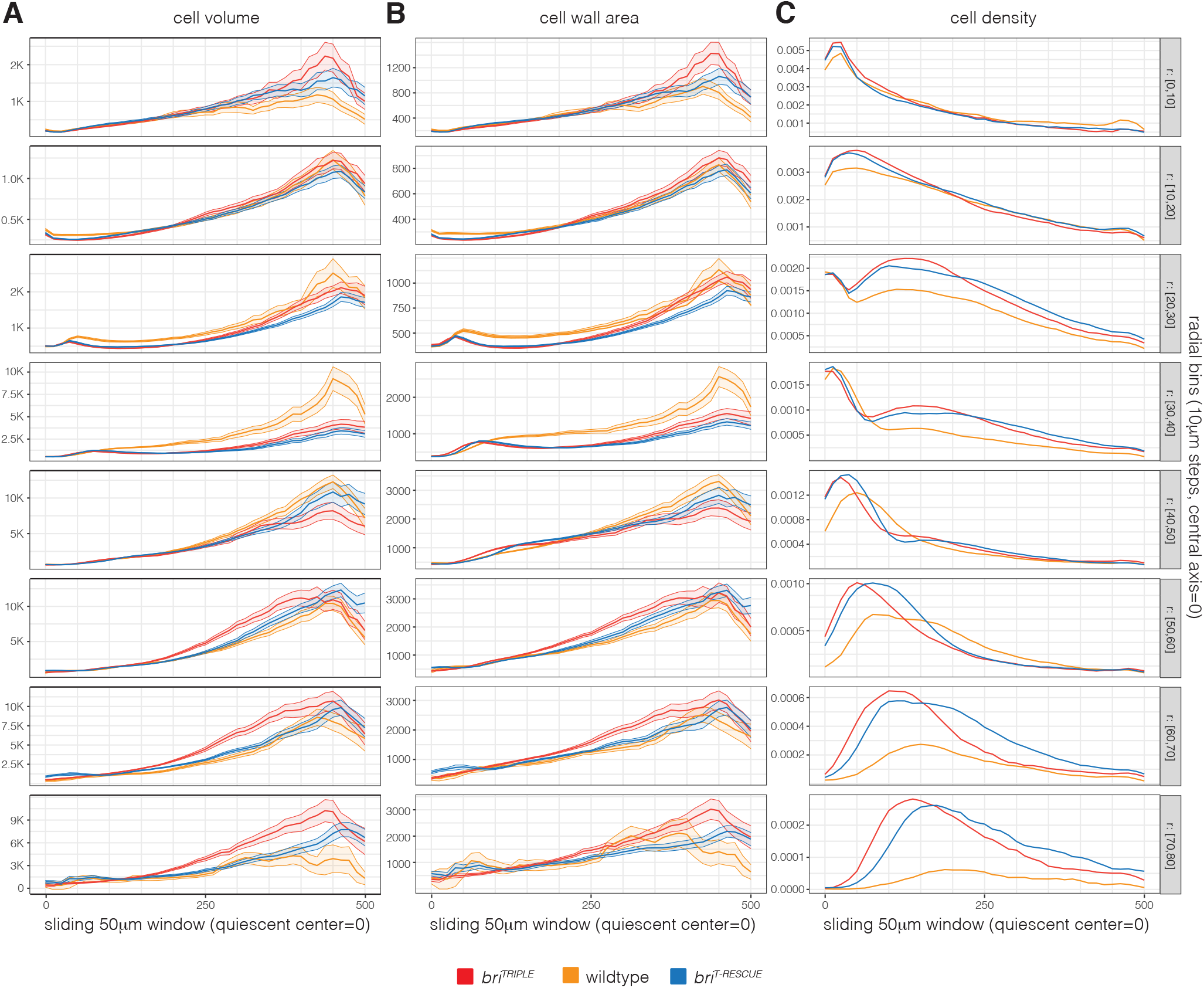
Comparative longitudinal sliding window analysis of standardized root models 2. Continuation of the analysis presented in Figure S6, indicating cell volume (*μ*m^3^) (A), cell wall area (*μ*m^2^) (B), and cell density (C) for 11-12 roots per genotype combined. Shaded regions indicate the +/− standard error of the mean where applicable. sNote that curves within single panels are not always representing similar tissues, because of the wider stele of *bri^TRIPLE^* meristems.

**Figure S8.**
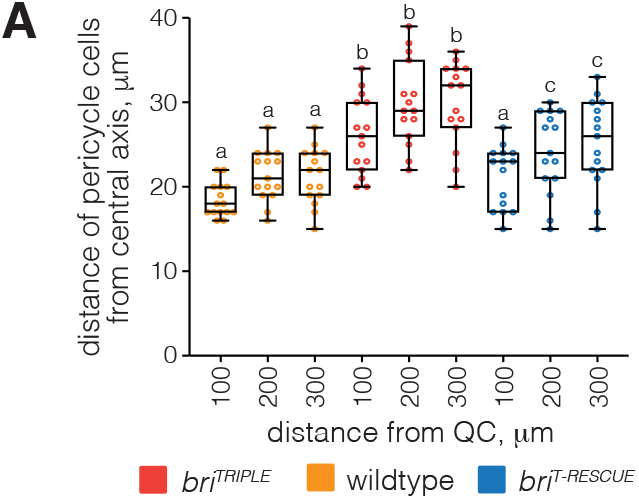
Stele width in wildtype, *bri^TRIPLE^* and *bri^T-RESCUE^* root meristems. (A) Average distance of pericycle cells from the central axis, for selected distances from the QC. Box plots display 2nd and 3rd quartiles and the median, bars indicate maximum and minimum. Statistical significance was determined by ordinary one-way ANOVA. Statistically significant different groups are indicated by different lowercase letters.

**Figure S9.**
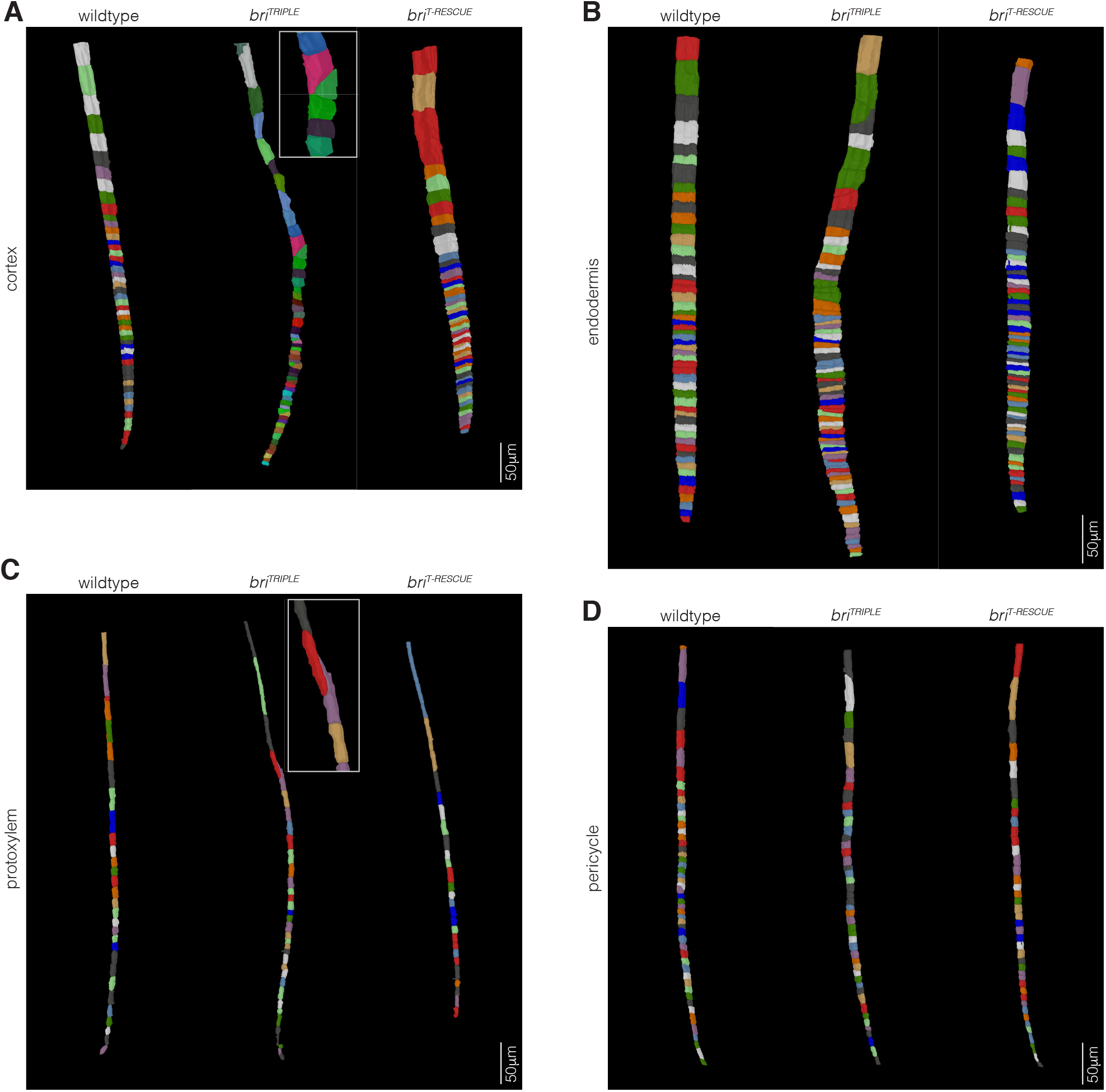
Examples of cell files in wildtype, *bri^TRIPLE^* and *bri^T-RESCUE^* root meristems. (A-D) Examples of isolated cell files for cortex (A), endodermis (B), protoxylem (C), and pericycle (D), obtained from segmentations via the *PlantSeg-MorphoGraphX-3DCellAtlas* pipeline. Note the frequently oblique cell division planes in *bri^TRIPLE^* mutants and the “ballooning effect”, highlighted by inlays.

**Figure S10.**
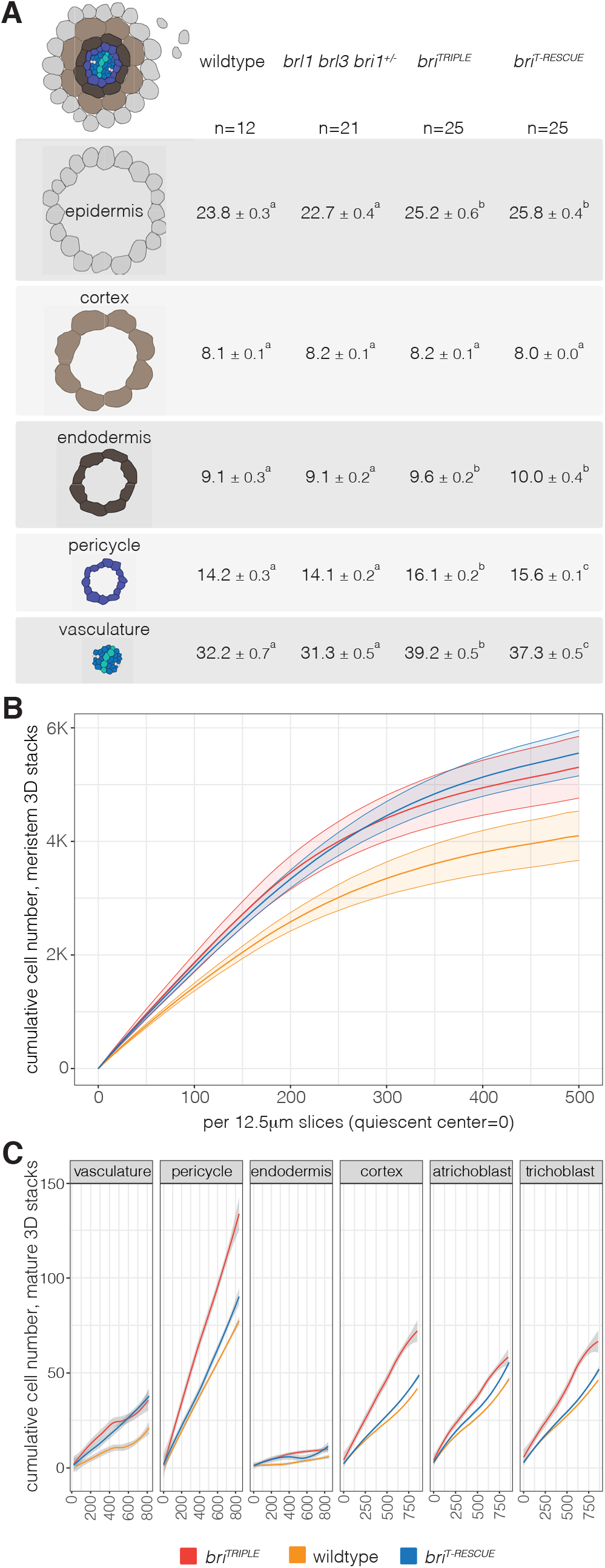
Quantification of cell files in wildtype, *bri^TRIPLE^* and *bri^T-RESCUE^* roots. (A) Quantification of cell file numbers in the different root tissues, obtained from histological cross sections taken at the level of differentiated protoxylem. Statistically significantly different groups are indicated by lower case letters. (B) Cumulative cell file number in root meristems, counted in 12.5 *μ*m slices from the QC to the elongation-differentiation zone, with +/− standard error of the mean. (C) Cumulative cell number in mature root area segmentation spanning 800 *μ*m in length and starting ~1 cm above the root meristem, with +/− standard error of the mean.

**Figure S11.**
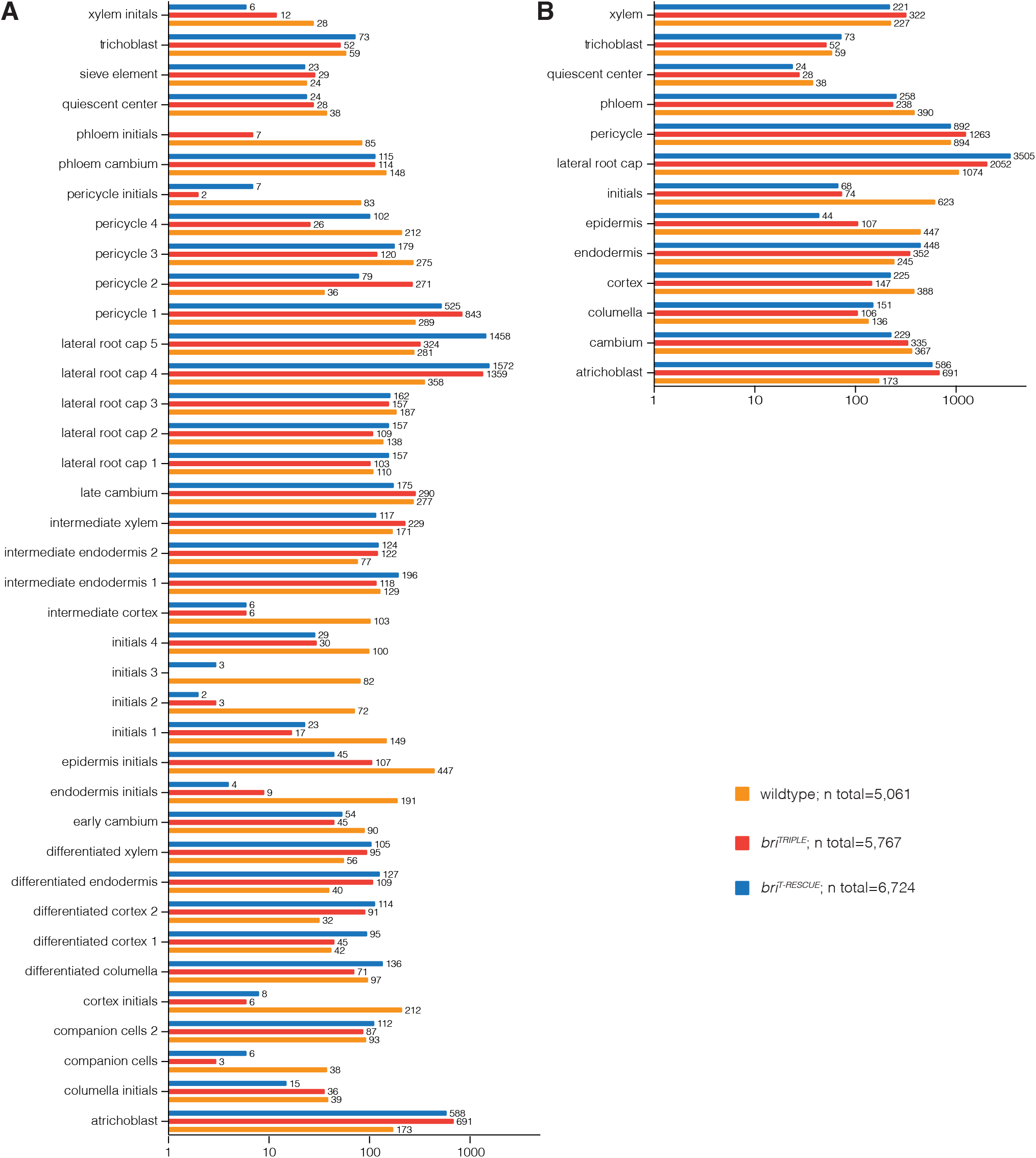
Cell type abundance in the scRNAseq datasets. (A) Absolute cell numbers per cluster in wildtype, *bri^TRIPLE^*, and *bri^T-RESCUE^* root meristem single cell transcriptomes, based on identity assignment with cell type-specific and stage-specific marker genes (Wendrich et al., 2020) (see Tables S1-S3). (B) As in (A), for the 13 principal cell types (see Table S4).

**Figure S12.**
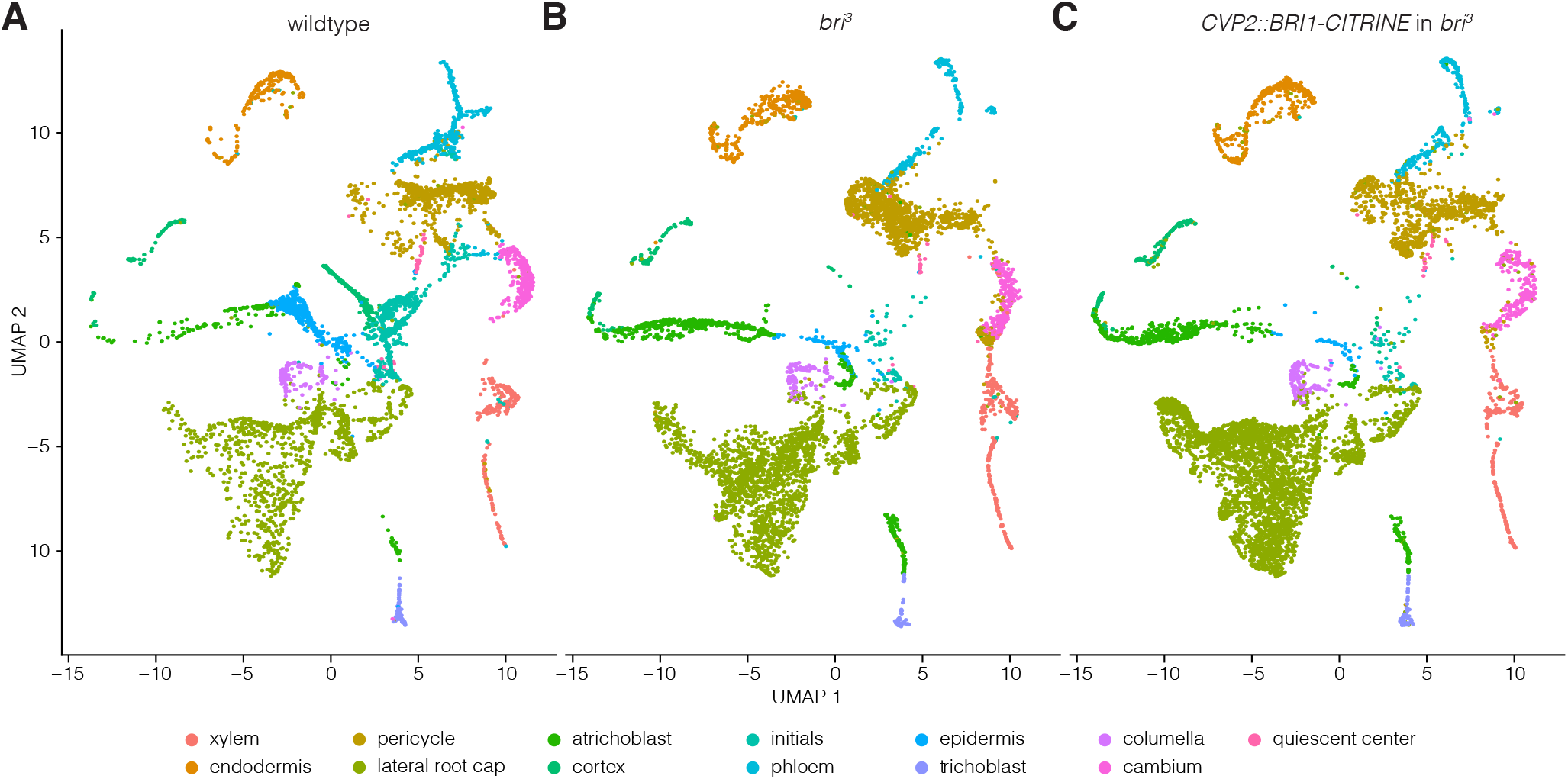
Overview of scRNAseq cell type clusters. (A-C) Uniform Manifold Approximation and Projection (UMAP) of wildtype (A), *bri^TRIPLE^* (B), and *bri^T-RESCUE^* (C) single cell transcriptomes, clustered based on assigned principal cell identities established by cell type-specific marker genes (Wendrich et al., 2020).

**Figure S13.**
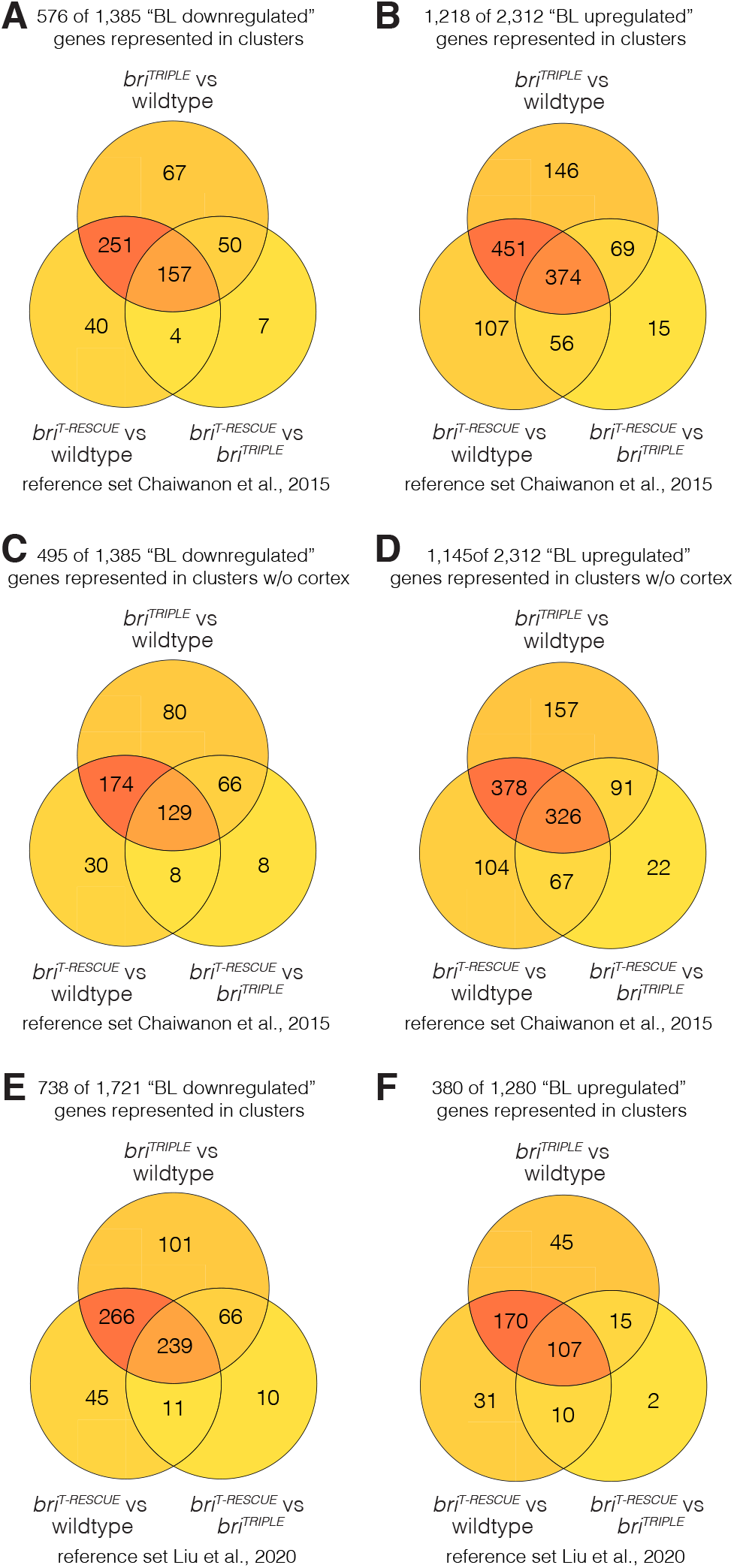
Comparison of scRNAseq data with known brassinosteroid-responsive genes. (A-B) Venn diagram illustrating the overlap between genes that were differentially expressed between genotypes in the scRNA-seq dataset, and genes that were found to be downregulated (A) or upregulated (B) in response to brassinolide treatment in the root (Chaiwanon and Wang, 2015) (see also Table S5 and Table S6). (C-D) As in (A-B), but with the cortex cells removed from the scRNA-seq dataset. (E-F) As in (A-B), for a different reference set of genes that are considered high confidence brassinosteroid-downregulated (E) or upregulated (F) (Liu et al., 2020).

**Figure S14.**
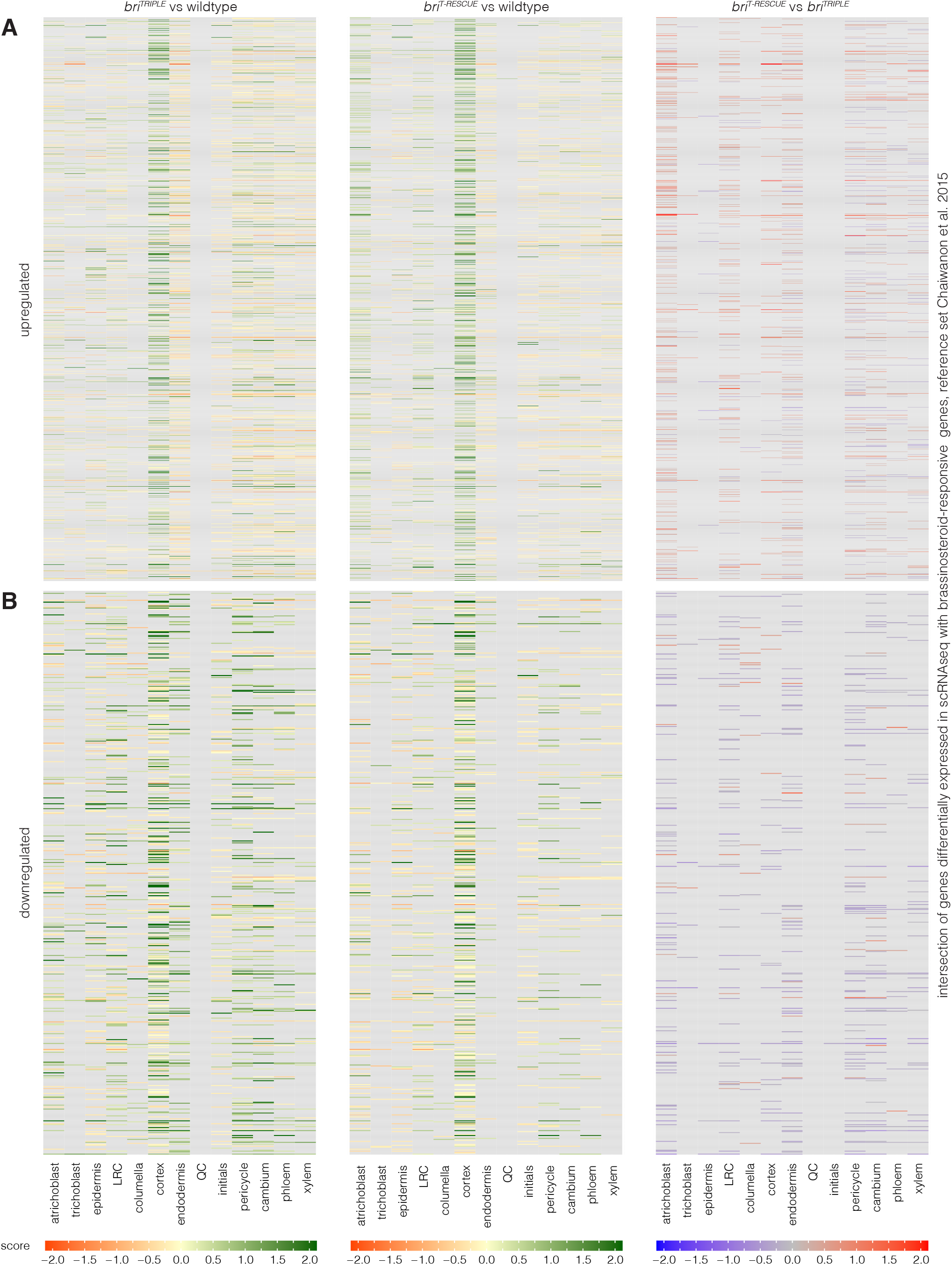
Brassinosteroid response in the scRNA-seq profiles. (A-B) Heatmap representing the expression of genes that were found to be upregulated (A) or downregulated (B) in response to brassinolide treatment in the root (Chaiwanon and Wang, 2015). The scRNA-seq dataset (Table S4) comprised 576 of 1,385 downregulated (Table S5) and 1,218 of 2,312 upregulated (Table S6) genes. Colour scales with a fixed range from −2 to 2 indicate the expression score (logFC x percentage of cells expressing gene in cluster X in genotype a / percentage of cells expressing gene in cluster X in genotype b). Genes that were not detected in the cluster or had an adjusted *p* value >0.05 are displayed in grey.

**Figure S15.**
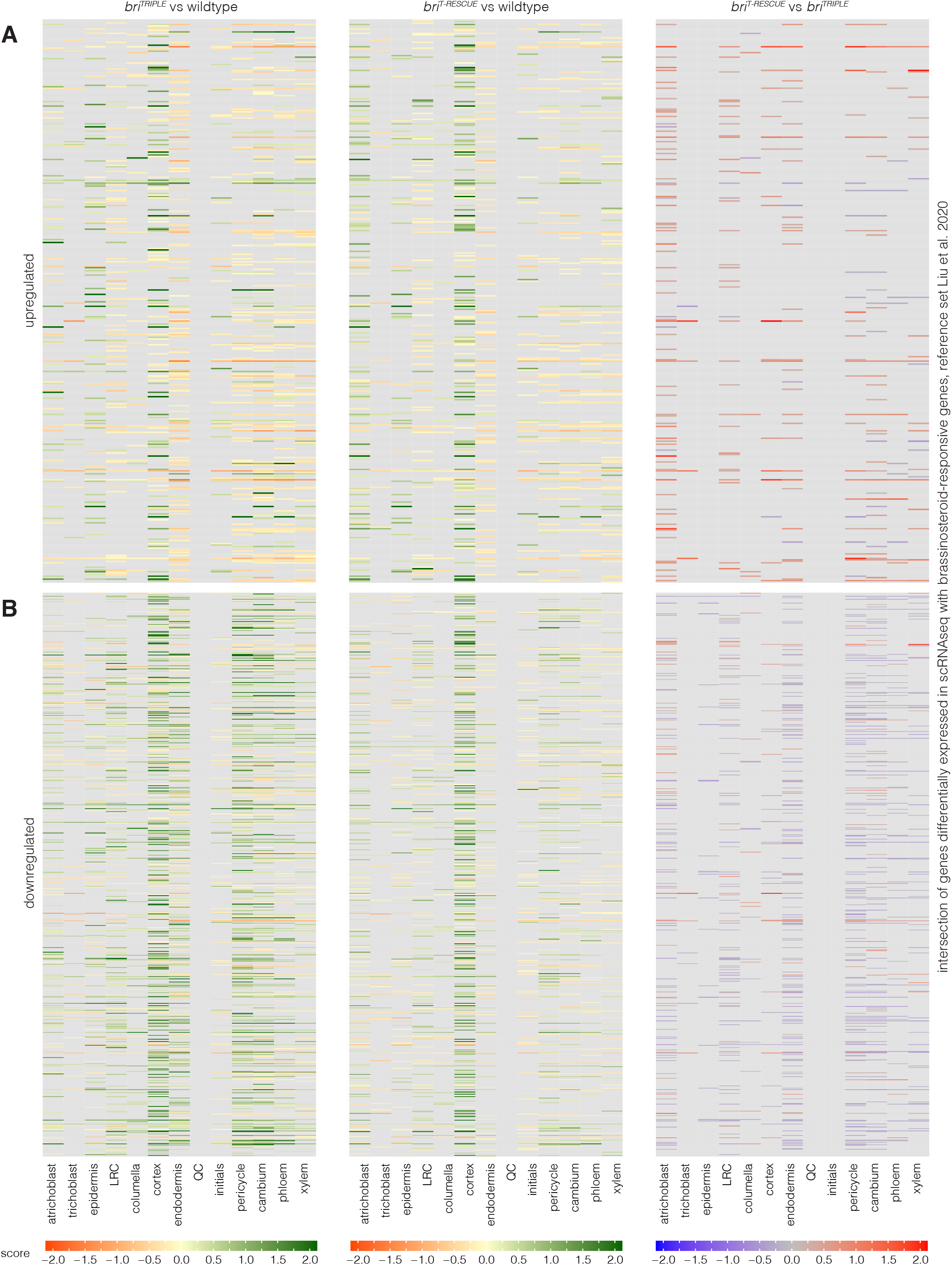
Brassinosteroid response in the scRNA-seq profiles. (A-B) Heatmap representing the expression of genes that are considered high confidence brassinosteroid-upregulated (A) or downregulated (B) (Liu et al., 2020). The scRNA-seq dataset comprised 380 of 1,280 upregulated and 738 of 1,721 downregulated genes. Colour scales with a fixed range from −2 to 2 indicate the expression score (logFC x percentage of cells expressing gene in cluster X in genotype a / percentage of cells expressing gene in cluster X in genotype b). Genes that were not detected in the cluster or had an adjusted *p* value >0.05 are displayed in grey.

**Figure S16.**
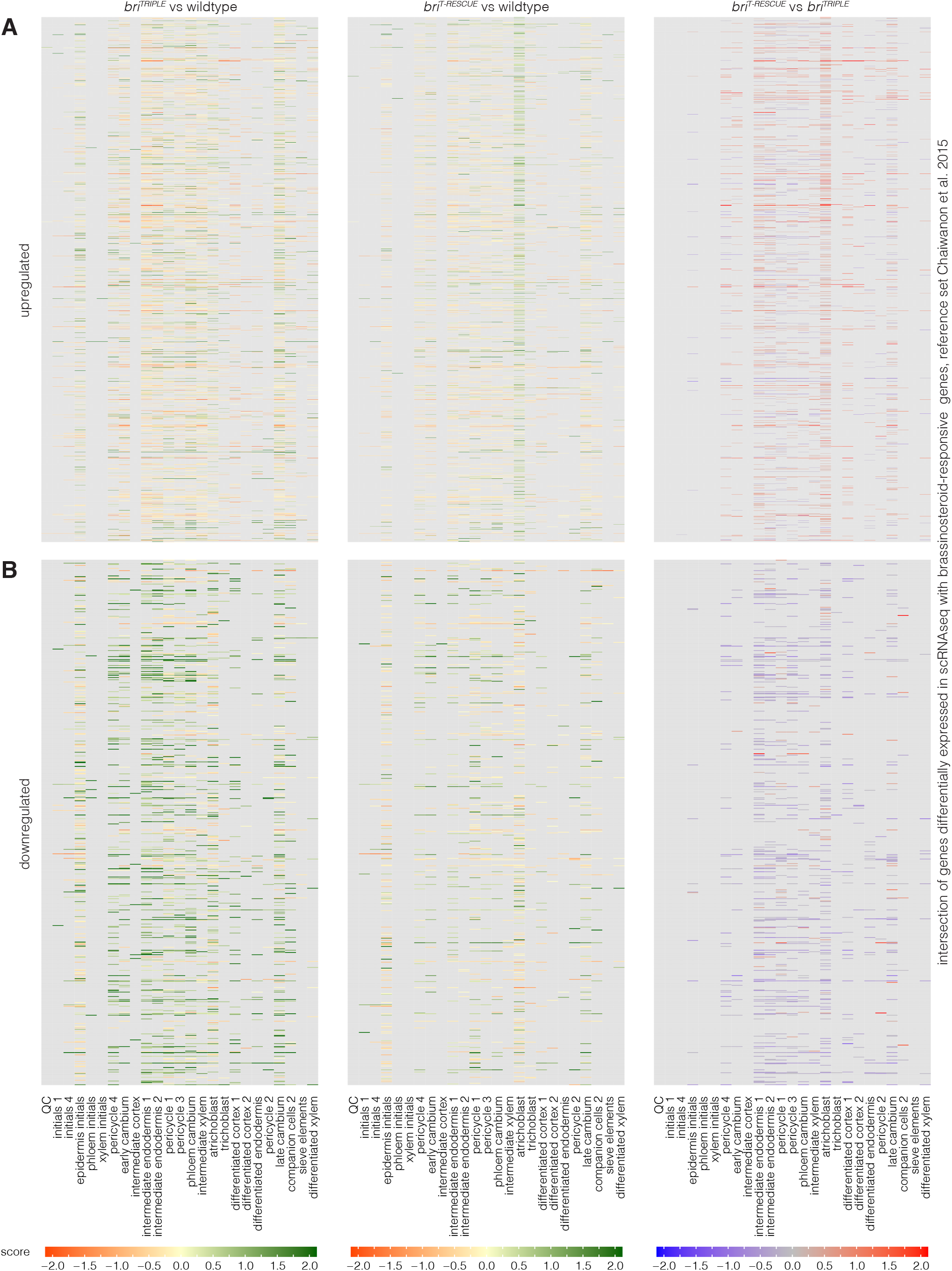
Spatial distribution of brassinosteroid response in the scRNA-seq profiles. (A) Heatmap representing the expression of genes that were found to be upregulated (A) or downregulated (B) in response to brassinolide treatment in the root (Chaiwanon and Wang, 2015), similar to Figure S14, but for subclusters and ordered with respect to stage-specific markers in increasing distance from the QC.

**Figure S17.**
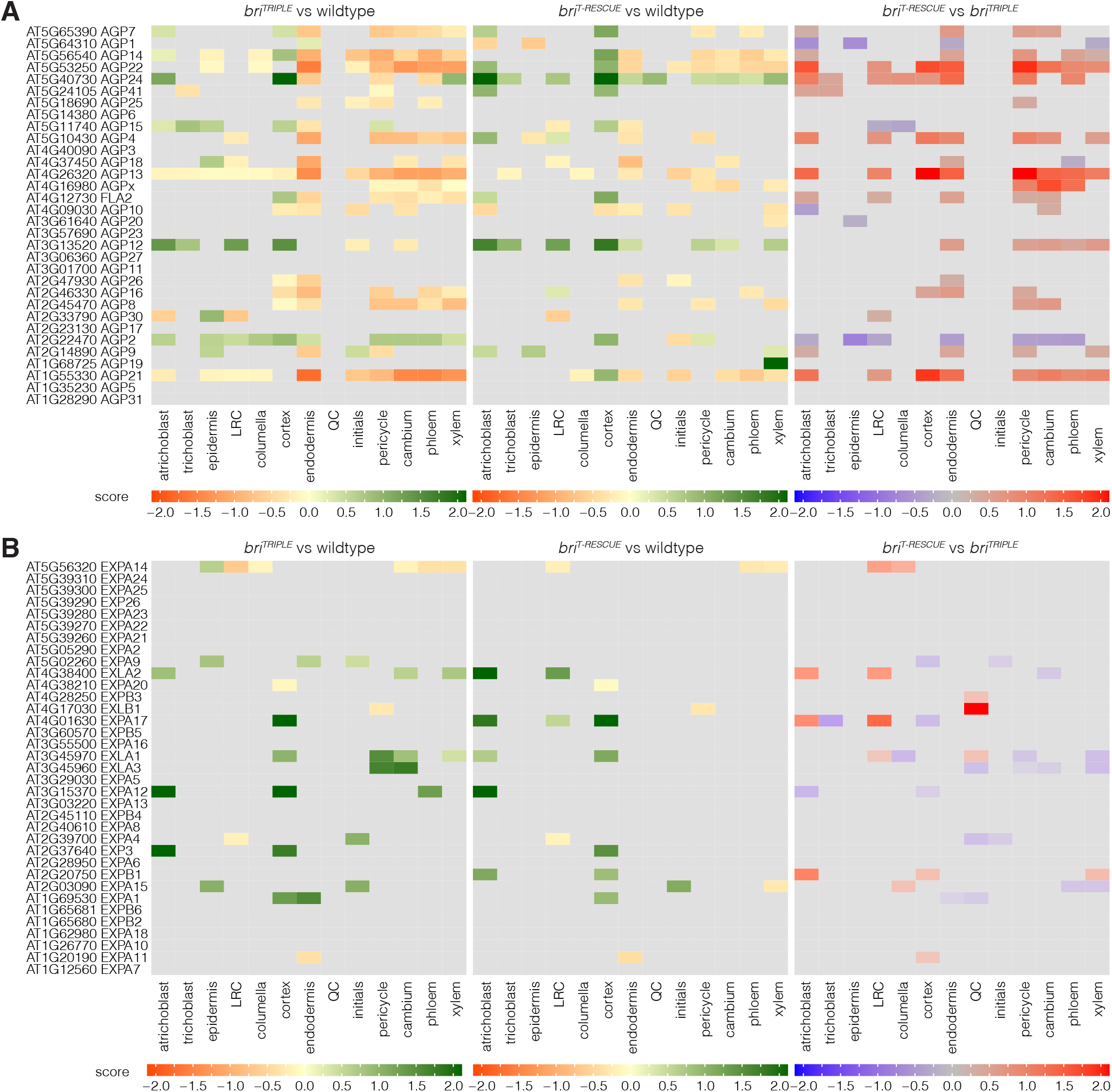
*AGP* gene expression in the scRNA-seq profiles. (A) Heatmap indicating the gene expression score of the 32 arabinogalactan proteins or peptides (AGPs) in the 13 general cell types. Note that AGPs show a consistent downregulation in the stele tissues and frequently also in ground tissues of *bri^TRIPLE^* meristems, which is largely normalized in *bri^T-RESCUE^* meristems. (B) Expansins are a gene family of comparable size to AGPs (35 genes in *Arabidopsis*) and are associated with cell expansion. Note that compared to AGPs, their expression is affected less frequently and expression profiles are more varied between the different genotypes.

